# Recombinant protein platform for high-throughput investigation of peptide-liposome interactions via fluorescence anisotropy depolarization

**DOI:** 10.1101/2025.05.12.653516

**Authors:** Antonis Margaritakis, Meirui Qian, David H. Johnson, Wade F. Zeno, Tobias S. Ulmer, Peter J. Chung

## Abstract

Many cytosolic proteins critical to membrane trafficking and function contain an unstructured domain that can bind to specific membranes, with a transition into an amphipathic helix induced upon membrane association. These inducible amphipathic helices often play a critical role in organelle recognition and subsequent function by these cytosolic proteins, but the tools and techniques used to characterize affinity towards specific membranes are low-throughput and highly dependent on the solubility of the inducible amphipathic helix. Here, we introduce a modular recombinant protein platform for rapidly measuring the binding affinity of inducible amphipathic helices towards a variety of membrane compositions and curvatures using high-throughput fluorescence anisotropy measurements. Inducible amphipathic helices are solubilized with a fluorescently tagged small ubiquitin-like modifier (SUMO) protein and binding to membranes quantified by leveraging the unexpected decrease in fluorescence anisotropy upon binding, a phenomenon previously observed but not well understood. By using fluorescence anisotropy decay measurements and solution NMR experiments, we deduce that this phenomenon likely occurs due to the local increase in fluorophore motion upon binding to the membrane. Altogether, this recombinant protein platform can be readily applied to any inducible amphipathic helix of interest, allowing for detailed investigation of the specific membrane biochemical parameters facilitating binding.

## Introduction

Inducible amphipathic helices are protein motifs that transition from a disordered state in solution to an amphipathic helix upon association with a particular membrane. These motifs are often found in proteins related to membrane remodeling, signaling, and trafficking^1–7^, with their peptide sequence driving their ability to recognize distinct membranes. For example, helix 0 of Amphiphysin (part of the NBAR superfamily of proteins) directly influences the protein’s ability to associate with membranes, with its presence linked to reduced curvature sensitivity and weaker binding^8–10^. The N-terminal domain of the Endophilin family of proteins has an increased affinity for lipids found in the mitochondria and endosomes, consistent with its putative biological function^11–13^. Similarly, the N-terminal domain of Huntingtin binds preferentially to membranes with high anionic lipid content and low cholesterol, mimicking mitochondrial membranes^14^. The ability of the inducible amphipathic helix to detect and bind to the correct cognate membrane is critical for proper function of the parent protein^9,15^.

Although inducible amphipathic helices are crucial for subcellular localization, systematically identifying the parameters that govern their lipid targeting remains experimentally challenging^16^. First, their ability to bind and insert into membranes necessitates a degree of hydrophobicity that, without its linkage to the necessarily more soluble parent protein, leads to instability in solution^17^. For example, some isolated N-terminal domains of the endophilin isoforms exhibit poor solubility and readily precipitate out of solution^13,18^. Furthermore, common techniques for extracting binding isotherms (such as tryptophan fluorescence, isothermal titration calorimetry, circular dichroism, etc.) are inherently low throughput and sample intensive, with a single isotherm for one lipid composition potentially requiring hours to measure and large amounts of protein and lipid. Additionally, the sheer diversity of lipid species within trafficking organelles creates an enormous compositional phase space, making comprehensive exploration of the determinants of selective membrane binding a daunting experimental challenge^19^. Understanding the underlying parameters that govern this binding behavior demands a highly parallelized and systematic approach, as comparisons across studies of individual proteins are often challenging and often lead to contradicting conclusions. For instance, both the curvature inducing abilities (or absence thereof) of the helix 0 of Amphiphysin^11,18,20,21^ and the role of the helix 0 of N-BAR Endophilin in membrane curvature generation^18,22,23^ have been difficult to determine.

Here, we introduce a modular recombinant protein platform for high-throughput quantification of the binding affinity of inducible amphipathic helices to model membranes using fluorescence anisotropy measurements, commonly available in microplate readers. Our system solubilizes the peptide of interest using a C-terminally linked SUMO (Small Ubiquitin-like Modifier) protein (Uniprot accession #: Q12306). Not only does the SUMO protein helps solubilize our peptide of interest, it greatly amplifies expression yields in E.coli^24^. Moreover, our platform is co-expressed in an N-terminal acetylation system to better reflect modifications found in their native environments^25^. A fluorescent label (unless otherwise stated, an Oregon Green 488 conjugated through a maleimide reaction to a cysteine mutated into the C-terminal domain) allows for spectroscopic quantification of the peptide-liposome interaction via fluorescence anisotropy (Figure 1a) and a downstream TEV-cleavable hexahistidine tag facilitates purification. Strikingly, we detect a decrease in fluorescence anisotropy upon binding (Figure 1b). We expected the apparent larger size of the bound platform/vesicle system (upon inducible amphipathic helix binding) to lead to elevated fluorescence anisotropy, consistent with the slower rotational diffusion of the bound state^26^. The observed anisotropy decrease indicates a counterintuitive increase in rotational diffusion of our recombinant platform when bound to the much larger vesicle. However, given the clear sigmoidal characteristic of the anisotropy signal at higher lipid to protein ratios (albeit, in an unexpected direction), we continued to investigate the binding behavior using a complementary technique, tryptophan fluorescence. Surprisingly, we found strong quantitative agreement between our fluorescence anisotropy and tryptophan fluorescence measurements, suggesting that the decrease in fluorescence anisotropy corresponded to the bound state of our recombinant platform. To illuminate the mechanism that leads to this phenomenon, we investigated the dynamic structure of our recombinant platform using a combination of fluorescence anisotropy decay and nuclear magnetic resonance measurements. These techniques reveal the presence of a secondary motion (stemming from the disordered nature of the fluorescently-tagged C-terminal domain) that conformationally relaxes and allows for an increase in the local mobility of the fluorophore upon membrane binding. This increase in local mobility dominates the overall rotational motion of the bound platform/vesicle system, leading to an observed reduction in the steady state fluorescence anisotropy. The robustness of this phenomenon allows for this platform to be readily applied as a high-throughput screening tool to determine the dynamic parameters governing peptide-membrane interactions. By systematically mapping the regions of membrane phase space where protein binding is maximized, our platform enables a rigorous investigation of the peptide code responsible for precise protein subcellular localization, paving the way towards reconciling peptide sequence with organelle specificity.

**Figure 1.**
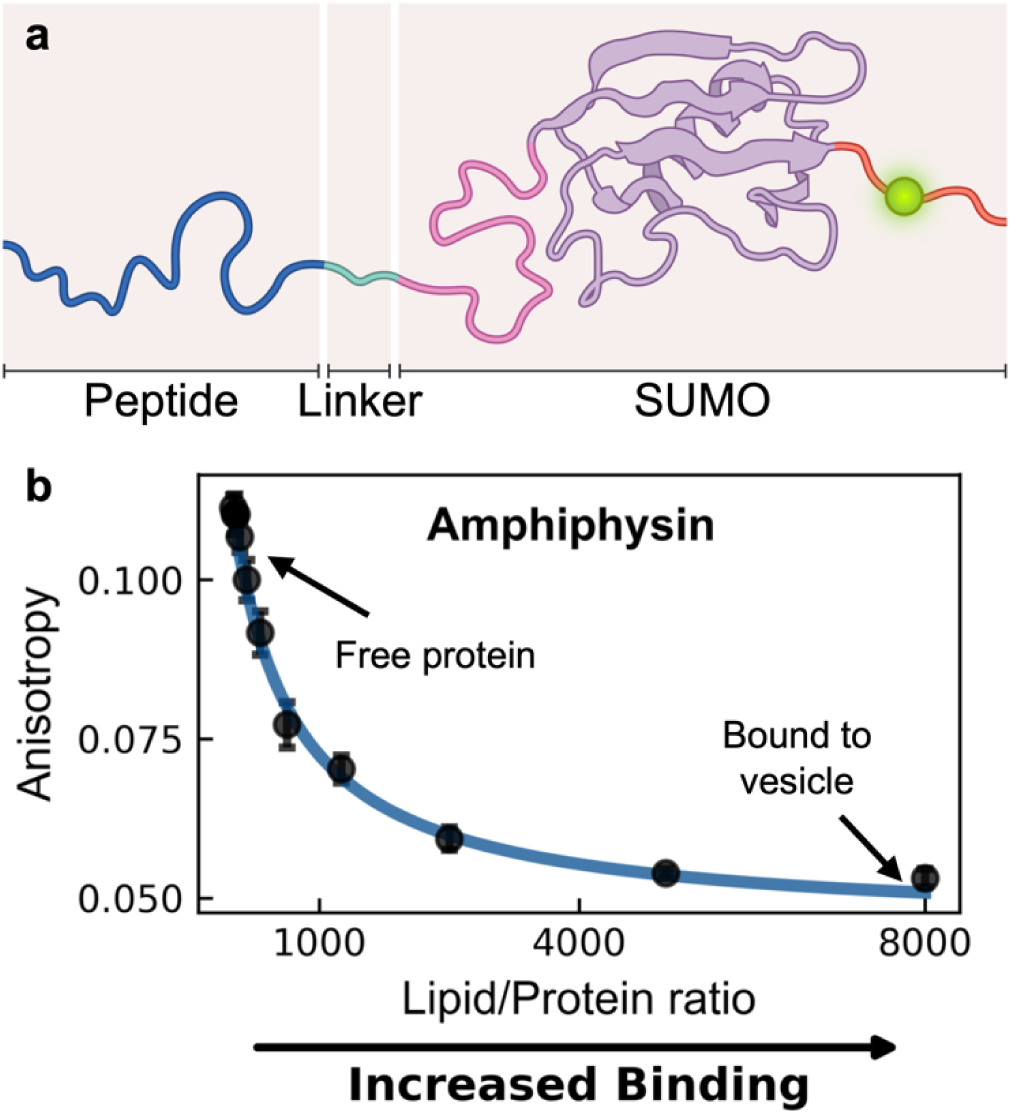
Our recombinant protein platform exhibits a decrease in fluorescence anisotropy upon binding to a vesicle. (a) Schematic representation of the SUMO protein platform. Peptides of interest (dark blue) are engineered at the N-terminus, linked to SUMO via soluble linker (light blue) and a mutated cysteine to facilitate fluorophore attachment (green). A cleavable C-terminal hexahistidine tag (not depicted) facilitates purification and cleaved prior to usage. (b) An unexpected decrease in fluorescence anisotropy is observed when measuring the N-terminal domain of Amphiphysin engineering to our platform upon increasing titration of 67 nm diameter DOPC/DOPS (70/30) vesicles. Data was fit using a membrane partitioning model (blue) as described in Methods.

### High-throughput detection of peptide-liposome binding

To leverage the decrease in fluorescence anisotropy of our recombinant platform as a high-throughput probe for binding affinity, we constructed binding isotherms for two distinct peptides against a 4 x 3 array of distinct vesicles. Along one axis of the array, we increased 1,2-dioleoyl-sn-glycero-3-phosphoethanolamine (DOPE) content (against decreasing 1,2-dioleoyl-sn-glycero-3-phosphocholine [DOPC] background and constant 30% 1,2-dioleoyl-sn-glycero-3-phospho-L-serine [DOPS]). On the other axis, we increased membrane curvature by extruding vesicles with decreasing diameters (Figure 2a). We chose DOPE as its smaller headgroup relative to DOPC and DOPS promotes surface defects that may allow for an easier amphipathic helix insertion^3,27^. Similarly, many inducible amphipathic helices preferably associate with highly curved surfaces^28^ (or even promote high curvatures)^29,30^, potentially due to a curvature induced bilayer asymmetry facilitating easier access to the hydrophobic acyl chain region.

**Figure 2.**
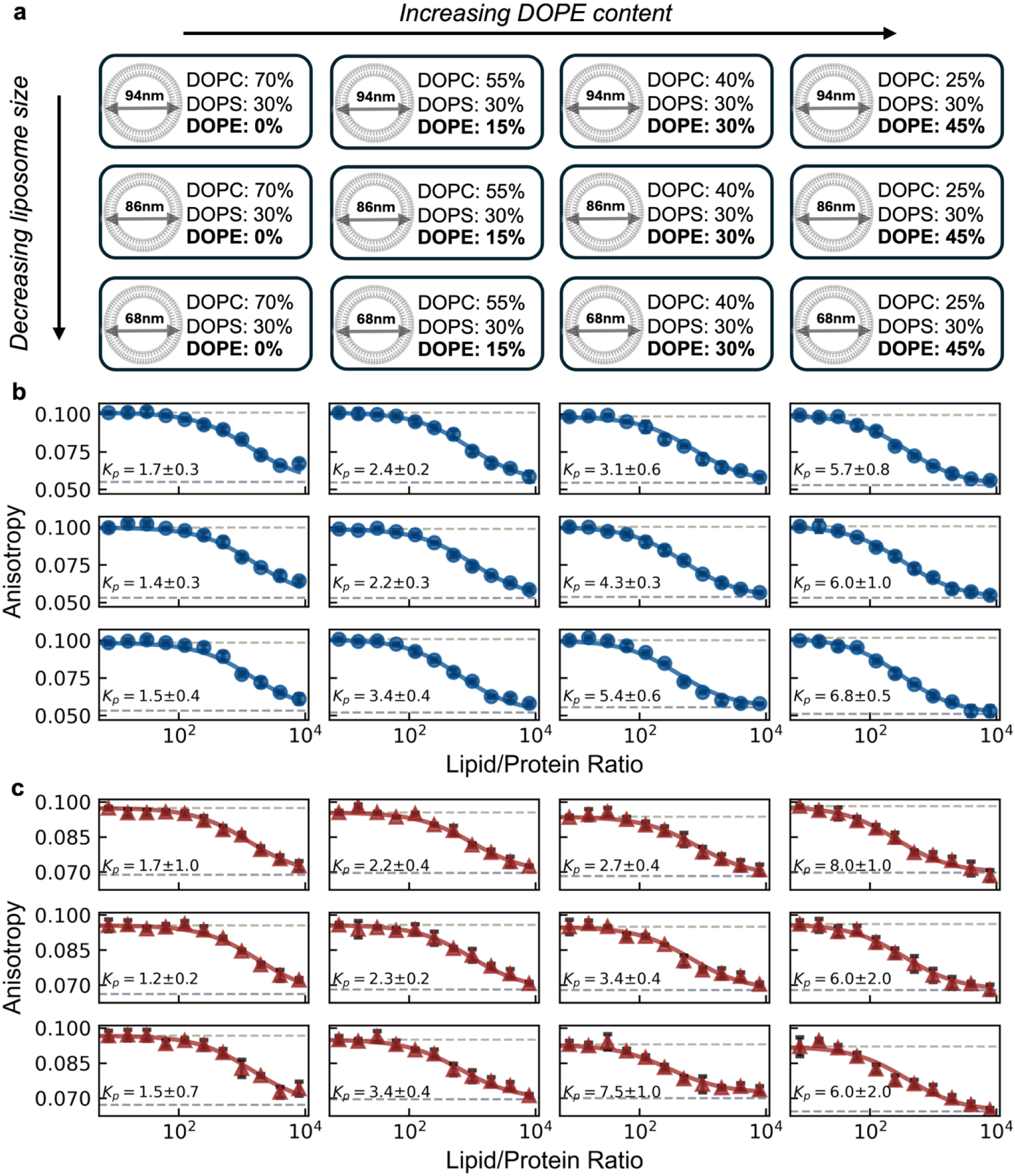
**High-throughput detection of inducible amphipathic helix binding to liposomes is possible**. (a) Schematic representation of the vesicle array used to assess protein binding in different membrane composition and curvatures. Four vesicle composition batches were prepared: DOPC/DOPS/DOPE = (70-X)/30/X with X = (0, 15, 30, 45), each extruded to three different final diameters D ≈ (94 nm, 86 nm, 68 nm). **(b, c)** Fluorescence anisotropy measurements of our recombinant platform engineered with the inducible amphipathic helices of **(b)** CHMP4B (AA:1-19, blue circles) and **(c)** Huntingtin (AA:1-17, red triangles) with increasing vesicle concentration. Data points show the mean of 3 technical replicate measurements for each lipid/protein ratio, with standard deviations plotted (black error bars). Data were fit using a membrane partitioning model (solid lines) with partition coefficient values shown as x10^5^.

We expressed and purified our recombinant platforms presenting peptides corresponding to the inducible amphipathic helices (AH) of the CHMP4B (AA1-19, Uniprot: Q9H444) subunit from the ESCRTIII protein complex and Huntingtin (AA1-17, Uniprot: P42858), hereinafter referred to as the CHMP4B AH and Htt17 AH respectively. We constructed each binding isotherm by measuring the fluorescence anisotropy at 12 lipid/protein ratios of our 4 x 3 array. Each 100 μL sample had a protein concentration of 0.25 μM, with three technical replicates for each lipid/protein ratio. The resulting binding isotherms were fit to the partition equilibrium function assuming a single bound state (see Methods), yielding a corresponding partition coefficients (Figure 2b,c, Figure S1). The CHMP4B AH exhibited a considerable preference for membranes with higher DOPE content across the range of membrane curvatures tested. Irrespective of curvature, the fitted partition coefficient increases ∼5-fold for higher DOPE content. Interestingly, the CHMP4B AH exhibited a preference for higher membrane curvatures at higher DOPE content (30%, 45%), suggesting synergistic effects. A similar behavior is observed for the Htt17 AH, which showed increased partitioning towards membranes with higher DOPE content. Vesicles with 15% and 30% DOPE content appeared to facilitate the emergence of a curvature preference in binding, with 68 nm diameter vesicles exhibiting a significantly higher binding affinity than vesicles with diameters 86 nm and 94 nm.

### Tryptophan fluorescence measurements validate our recombinant platform as a tool for measuring binding affinity

Having extracted partition coefficients from our recombinant platform in a high-throughput manner, we sought to rigorously compare our probe to values derived from a complementary technique. Although slower and much more sample intensive, tryptophan fluorescence is considered a rigorous, quantitative technique which can evaluate the binding of inducible amphipathic helices with membranes^31–33^. It leverages the natural fluorescence of tryptophan which changes depending on the polarity of its immediate environment (with an emission centered ∼350 nm in aqueous solvents and ∼330 nm when submerged in the nonpolar acyl chain layer). Conveniently, all peptides tested contain at least one phenylalanine, which can be mutated into a physiochemically similar tryptophan, enabling parallel measurements using fluorescence anisotropy (Figure S2) and tryptophan fluorescence (Figure S3). We extracted binding isotherms of four distinct peptides to vesicles with systematically varied membrane compositions, specifically targeting compositions known to influence the binding affinity (DOPE and cardiolipin content for Endophilin-B1^3,13^ and Huntingtin^3,14^, respectively) or suspected of doing so (cardiolipin and cholesterol content for Amphiphysin^34,35^ and CHMP4B^4,36^, respectively) (Figure 3). Binding isotherms showed strong quantitative agreement between fluorescence anisotropy and tryptophan fluorescence measurements, with consistent results across systematically varied membrane compositions. We also performed control measurements using a platform without conjugated peptides to confirm that the binding isotherms exclusively reflect interactions with the inducible amphipathic helices (Figure S4).

**Figure 3.**
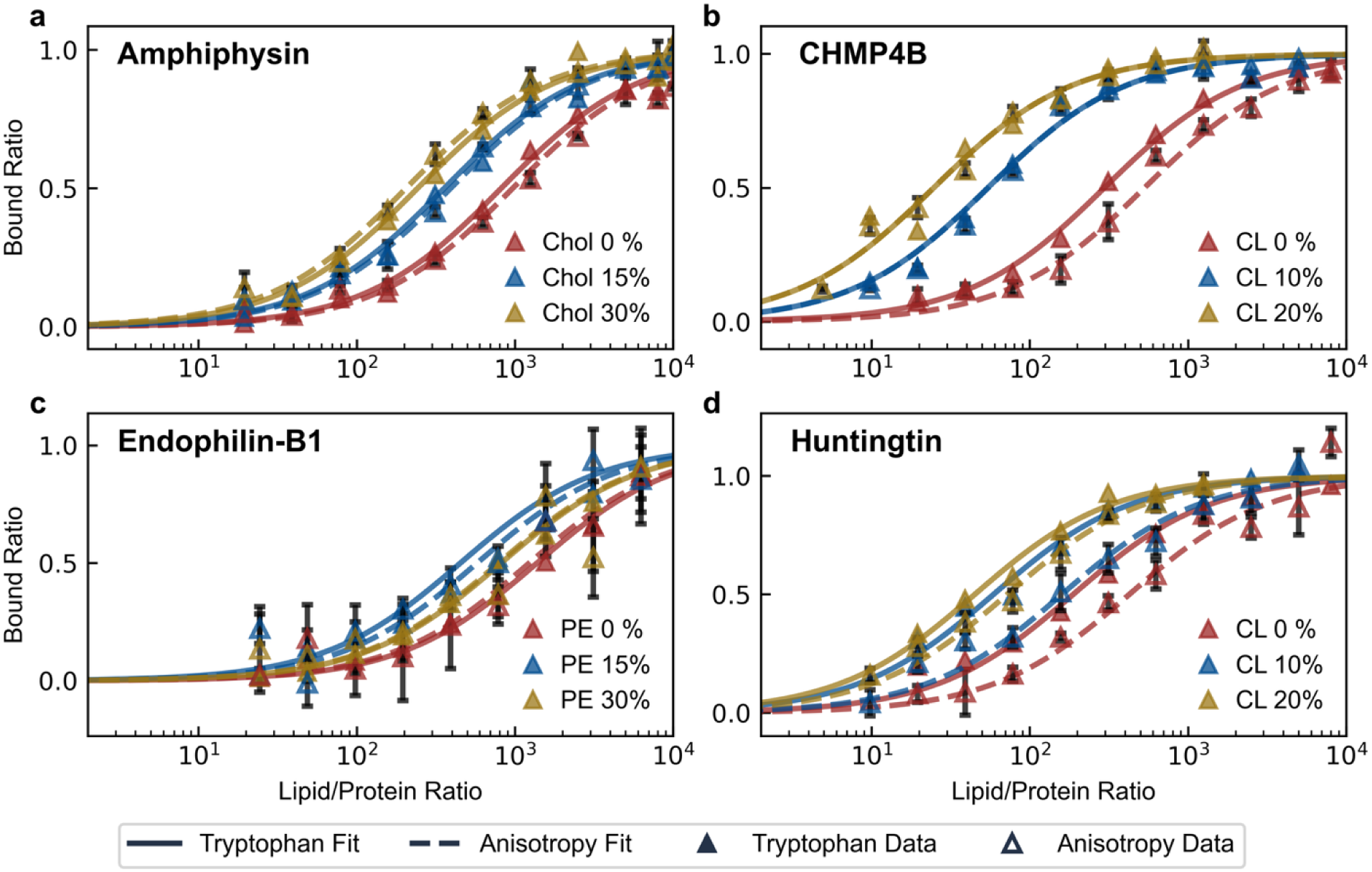
Binding isotherms derived from fluorescence anisotropy and tryptophan fluorescence show strong quantitative agreement. Parallel fluorescence anisotropy and tryptophan fluorescence measurements were taken with our recombinant platform engineering with the inducible amphipathic helices of **(a)** Amphiphysin (AA: 1-25, F9W mutant), **(b)** CHMP4B (AA: 1-19, F8W mutant), **(c)** Endophilin-B1 (AA: 1-33, F18W mutant) and **(d)** Huntingtin (AA: 1-17, F11W mutant). The binding of each platform was measured against changes in a variable lipid membrane content either known or suspected to influence binding affinity: cholesterol (Chol) for Amphiphysin, cardiolipin (CL) for CHMP4B, phosphatidylethanolamine (PE) for Endophilin-B1, and cardiolipin (CL) for Huntingtin. In all cases, vesicles were measured to be ∼70 nm diameter, composed of DOPC/DOPS/variable lipid at (70-X)/30/X molar ratios.

During these measurements, we uncovered a previously unreported result. We detect a significant monotonic increase in partitioning for helix 0 of Amphiphysin towards vesicles containing increased cholesterol content. Helix 0 of Amphiphysin exhibits ∼2-fold and ∼4-fold higher partition coefficients for vesicles containing 15% and 30% cholesterol, respectively, compared to vesicles without cholesterol. In both cases, binding isotherms produced from the two techniques are in strong quantitative agreement (Figure S5).

In measuring the binding affinity of the CHMP4B AH, both techniques exhibited clear preferential partitioning into membranes with higher cardiolipin content. A 10% cardiolipin membrane content in vesicles yields a 5-fold increase in partitioning relative to vesicles containing none, while further addition of cardiolipin to a 20% ratio yields a further ∼3-fold increase. Notably, both techniques consistently captured the non-monotonic binding behavior of helix 0 of Endophilin-B1 in response to increasing DOPE concentrations (Figure S5). For vesicles composed of DOPC/DOPS (70/30) with an average diameter of ∼68 nm, we calculated a partition coefficient of (4±0.6)x10^5^, a result in close agreement with the value of (1.5±0.5)x10^5^ reported by Robustelli and Baumgart for vesicles similarly comprised of DOPC/DOPS (75/25) with an average diameter of 100 nm^13^.

Finally, we tested the Htt17 AH with vesicles containing increasing amounts of cardiolipin and both techniques unveiled a strong preference for membranes with higher cardiolipin content. Specifically, tryptophan fluorescence showed ∼2-fold and ∼3-fold increases for vesicles containing 10% and 20% cardiolipin, respectively, compared to vesicles with no cardiolipin content. In contrast, our platform’s fluorescence anisotropy decrease indicated ∼3-fold and ∼9-fold increases for the same compositions. While the magnitude of response differs between the two methods, both techniques consistently capture the same qualitative trend (namely, the preference of Htt17 AH for cardiolipin-containing membranes). We do not expect the binding isotherms between the two techniques to always be identical, as tryptophan fluorescence is a direct measure of binding while our platform’s fluorescence anisotropy reflects the proximity of the platform to the membrane. Nevertheless, we confirmed binding via the formation of an amphipathic helix (for all peptides) using circular dichroism spectroscopy (Figure S6). Our results clearly indicate that both methods can effectively discriminate the membrane compositional preferences of Htt17.

In all tested scenarios, the individual lipid preferences can be extracted from fluorescence anisotropy curves just as effectively as from tryptophan fluorescence data. The strong quantitative agreement between resultant binding isotherms and extracted partition coefficients underscores the robustness of our recombinant protein platform as a precision tool for measuring the binding affinity of inducible amphipathic helices, prompting further investigation into the mechanism by which a decrease in fluorescence anisotropy is observed upon binding.

### The fluorescence anisotropy decrease is dependent on apparent membrane charge, but not size or choice of fluorophore

To elucidate the mechanism by which our recombinant platform exhibited a decrease in fluorescence anisotropy upon binding, we devised a model version of our platform to systematically test the membrane conditions that could drive this effect. To achieve this, we replaced the peptide corresponding to an inducible amphipathic helix with hexahistidine. By incorporating a nickel-chelating lipid (10% 1,2-di-(9Z-octadecenoyl)-sn-glycero-3-[(N-(5-amino-1-carboxypentyl)iminodiacetic acid)succinyl]) into our vesicles, our model recombinant platform bound tightly to the membrane surface irrespective of specific peptide-vesicle interactions. This design enabled us to isolate the contribution of membrane itself to the decrease in fluorescence anisotropy, independent of the unique binding behavior of each peptide.

Using this model platform, we dissected the contribution of the following physiochemical parameters: vesicle size, membrane charge, salt conditions and fluorophore identity (Figure 4, Figure S7). These parameters were chosen as representative factors anticipated to most likely affect the local mobility of the negatively charged fluorophore (at physiological pH) within its immediate environment, potentially resulting in decreased fluorescence anisotropy when bound to the larger vesicle. The decrease in fluorescence anisotropy seemed to be independent of fluorophore and vesicle size (Figure S7, Figure 4a), yet highly dependent on membrane charge and salt concentration (Figure 4b,c), indicating an electrostatically driven phenomenon. Specifically, the progressive narrowing of the dynamic detection window with increasing salt concentration mirrored the effect exhibited with decreasing membrane negative charge. To ensure binding occurs under electrostatically neutral conditions, microscopy measurements confirmed colocalization occurs with electrostatically neutral DOPC vesicles (Figure S8). Taken together, these results strongly suggest that the decrease in fluorescence anisotropy exhibited by the system is strongly coupled to the amount of membrane charge “visible” to the fluorophore conjugated to the C-terminal domain of our platform. Supporting this hypothesis, altering the pH of our sample solution led to a decrease in fluorescence anisotropy for the free state of our recombinant platform, potentially due to pH induced charge changes in the fluorescently-labeled C-terminal domain of our platform (Figure S9). To further test this idea, we introduced a charge-altering mutation into the fluorescently-tagged C-terminal domain (R104H). This mutation narrowed the dynamic detection window without affecting the overall binding isotherm remained (Figure S10). These findings suggest that the decrease in fluorescence anisotropy is directly linked to membrane charge exposure and indicates that local conformational changes within the fluorescently-tagged C-terminal domain can modulate this effect without affecting overall binding.

**Figure 4.**
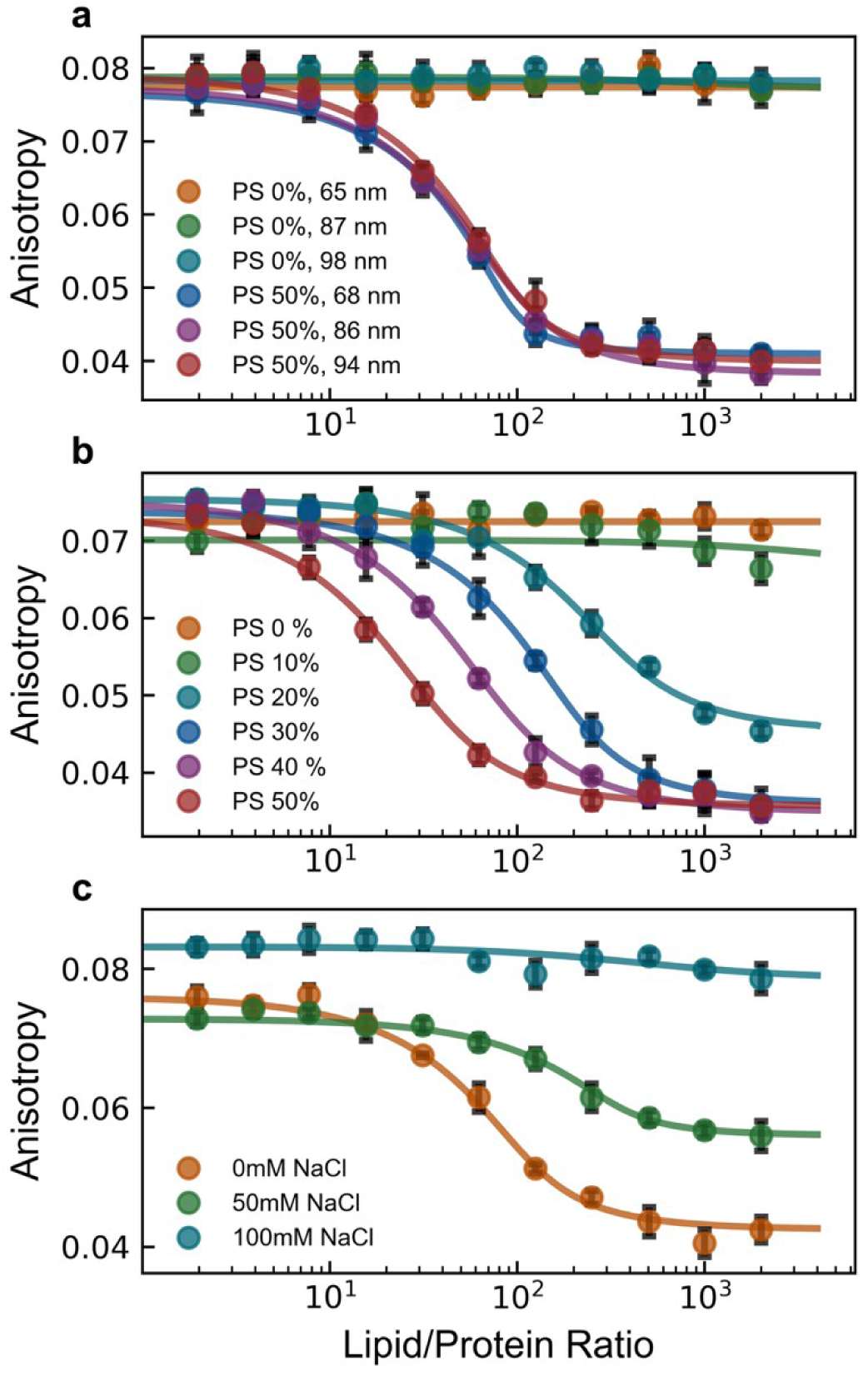
The decrease in fluorescence anisotropy is electrostatic in nature. Fluorescence anisotropy was measured for fluorescently-tagged hexahistidine model platform binding to nickel-chelating lipid containing vesicles against different conditions: **(a)** decreasing vesicle size (with lipid vesicle compositions of DOPC/DGS-NTA(Ni) = 90/10 or DOPC/DOPS/DGS-NTA(Ni) = 50/40/10), **(b)** membrane charge (with lipid vesicle compositions of DOPC/DOPS/DGS-NTA(Ni) = (90-X)/X/10 with X shown in the legend), and **(c)** buffer salt concentration (with lipid vesicle compositions of DOPC/DOPS/DGS-NTA(Ni) = 50/40/10). Data points are the mean of 4 replicate measurements for each lipid/protein ratio with standard deviations plotted (black error bars). Solid lines represent fits to a depletion model as described in methods. Unless otherwise noted, all vesicles were extruded to a final diameter of ∼95 nm.

### Decrease in fluorescence anisotropy occurs by increasing fluorophore mobility upon platform binding

To investigate the underlying conformational dynamics of our platform, we employed time-resolved fluorescence anisotropy decay. Unlike steady-state measurements, this technique delivers a precise excitation pulse of polarized light and monitors the time-dependent depolarization of the emitted fluorescence. Analyzing this decay over time allows us to separate the depolarization due to the potentially faster rotational diffusion of the fluorophore from the slower diffusion of the platform or platform/vesicle complex, providing mechanistic insights into the observed changes in steady-state anisotropy upon binding. These experiments required decay measurements in three states: free in solution, bound to negatively charged vesicles (to recapitulate the observed steady-state effect), and bound to neutral vesicles (which serves as a control, as it does not exhibit a decrease in anisotropy and thus reveals baseline behavior). We again used the hexahistidine model platform to examine the bound-state behavior. We first attempted to fit the decay of the three states with a model corresponding to a single rotational correlation lifetime, consistent with a fluorophore rigidly attached to the platform (in a free state) or the platform/vesicle complex (in the bound states). Unfortunately, this model fails to capture the early decay behavior of the fluorophore (Figure S11), suggesting a significant contribution from local fluorophore motion. As a result, we explored using a model in which there is an additional hindered degree of freedom, representing a fluorophore that is non-rigidly attached to the platform. This two-state hindered anisotropy decay model better represents fluorophores attached to macromolecules with segmented mobility^37^:

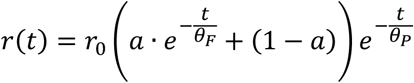

Where θ_*F*_ and θ_*P*_ represent the local and global rotational correlation times, respectively, and α is the fractional amplitude of the fast (local) motion in relation to the slower (global) motion. This model provides a better fit to the decay data, as indicated by smaller residuals and the absence of systematic deviations from baseline (Figure 5a, Figure S12a-f). For the free state, θ_*P*_ was determined to be (3±1) ns, closely matching the expected value of ∼4 ns calculated using the Stokes-Einstein-Debye equation. Samples containing vesicles yielded fitted parameters indicating a large global rotational correlation lifetime (θ_*P*_), several orders of magnitude greater than the local fluorophore rotational correlation lifetime (θ_*F*_) (Table S1). In this case, the fitted value of θ_*P*_ approaches infinity and indicated that the rotational diffusion is dominated by the vesicle, which has a calculated correlation time of ∼100μs. As a result, early time anisotropy decay is governed primarily by the local motion of the fluorophore, likely arising from its attachment to the disordered C-terminal domain of our platform. The fast local fluorophore motion, with a fitted θ_*F*_∼1ns (and significantly shorter than the fluorophore fluorescence lifetime of ∼4 ns), rapidly depolarizes the emitted photons and dominates the overall decay. We can further refine our bound state model to account for the slow platform/vesicle coupled motion by modifying the two-state decay in the limit where θ_*P*_ → ∞:

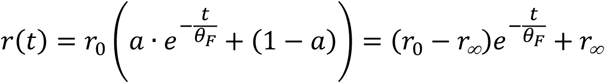

**Figure 5.**
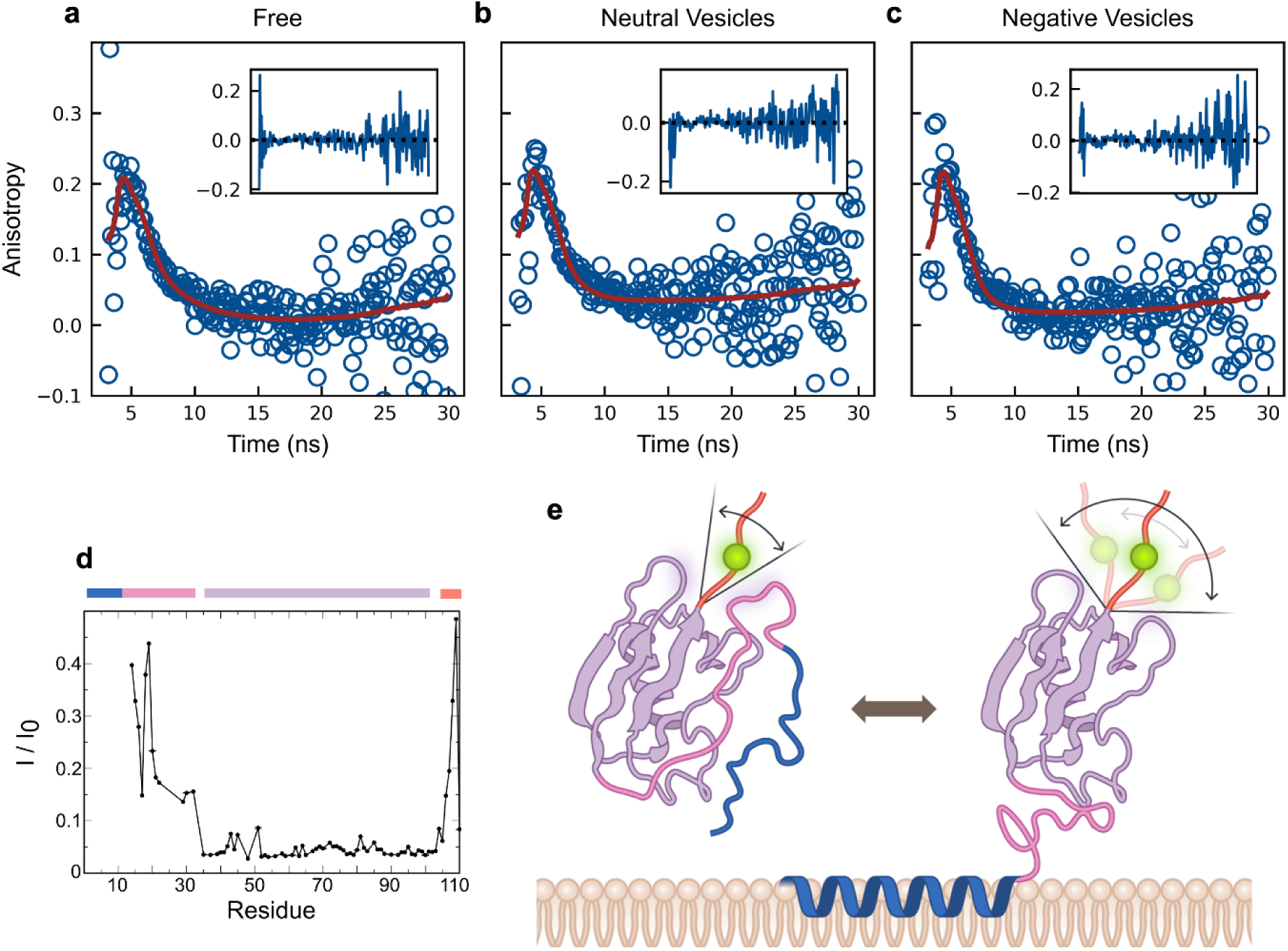
Fluorescence anisotropy decay and NMR measurements support a model by which decrease in fluorescence anisotropy occurs by a local increase in fluorophore mobility. Fluorescence anisotropy decay was measured for fluorescently-tagged hexahistidine model platform (25 nM) in 3 different states: **(a)** free in solution, **(b)** bound to neutral vesicles (DOPC/DGS-NTA(Ni) = 90/10, 25 μM) and **(c)** bound to negatively charged vesicles (DOPC/DOPS/DGS-NTA(Ni) = 50/40/10, 25 μM). Data were fitted as described in Methods using a two-state hindered anisotropy decay model for (a) and a hindered rotational diffusion model for (b) and (c). Insets show residuals of each fit. **(d)** Residual ^1^H^N^-^15^N signal intensities of the fluorescently-tagged hexahistidine model platform in the presence of negatively charged vesicles (DOPC/DOPS/DGS-NTA(Ni) = 50/40/10, 25 mM). For the protein remaining free in solution, three distinct areas of signal reductions were observed illustrating varying degrees of domain flexibility that are expected to be preserved in the vesicle-bound state. Vesicle-bound protein is tumbling too slowly to be observable. The differently affected areas are indicated by blue/pink, magenta and orange bars, and correspond to the protein domains shown in (e). **(e)** Schematic representation of our platform free in solution and bound to a vesicle, where the conformational freedom of the fluorescently-tagged domain increases.

This reduced two-state model (hindered rotational diffusion) provides a significantly improved fit to both the early and late anisotropy decay of the bound state (Figure 5b-c, Sup. S12g-l), successfully capturing system behavior in the presence of both negatively charged and neutral vesicles. The primary distinction between the decay behavior of the two vesicle species lies in the relative contribution of individual model parameters (*r_∞_* and θ_*F*_), which are modulated by the apparent presence or absence of membrane charge.

Taken together, the fluorescence decay data indicate the decrease in steady-state fluorescence anisotropy occurs because there is a transition. In the free state, there are meaningful contributions from both global and local motion of the platform and fluorophore, respectively. Upon binding, the overall decay is dominated by the local motion of the fluorophore and there is a concomitant increase in local mobility. We geometrically interpret this to represent a release of steric hindrance of the fluorophore. In the presence of a negatively charged membrane, the oscillatory local motion of the fluorophore increases due to the surface potential (an effect that does not occur with neutral membranes). The system’s sensitivity to variations in membrane charge and ionic strength further supports this mechanistic interpretation.

To corroborate our mechanistic interpretation that fluorophore mobility increases in the bound state of the platform, we investigated the structural and dynamic properties of our platform by nuclear magnetic resonance (NMR) spectroscopy. First, to verify that fluorophore interactions with the rest of the platform remained non-specific, we compared the NMR spectra of ^13^C/^15^N-labeled versions of our hexahistidine model platform with and without a fluorophore. Spectral differences predominantly localized to the C-terminal region where the fluorophore is conjugated (Figure S13), although some resonance signals within the SUMO domain were doubled indicating a non-negligible fluorophore proximity effect. Nevertheless, the small chemical shift differences between split resonances show that the structure of the folded SUMO domain remained virtually unchanged. Moreover, since specific SUMO-ligand interactions usually produce much larger shift changes^38^ and resonance doubling was also observed near the C-terminal conjugation site, we deem any SUMO-fluorophore interaction to be unspecific. Next, we investigated the NMR response of the platform in the presence of vesicles aiming to gain insight into the dynamic behavior of our platform. While the large particle size of the vesicle-bound platform did not allow the direct observation of the bound state by NMR, the protein fraction remaining free in solution experienced a relaxation contribution from its exchange with the vesicle-bound state. This contribution diminished the signal intensities of the disordered C-terminal region and the rest of the platform domains to different degrees (Fig. 5e) showing their dynamic, loose coupling. In the vesicle-bound state a loss of non-specific SUMO domain-fluorophore interactions and/or an altered proximity behavior may loosen this coupling even further. Overall, a lower steady state fluorescence anisotropy signal in the vesicle-bound relative to the free state is therefore compatible with an increase in the conformational freedom of the fluorescently-tagged C-terminal region of our platform upon vesicle binding.

### Conclusions/Further discussion

Leveraging an unexpected decrease in fluorescence anisotropy upon binding offers a powerful approach to studying molecular interactions. A similar phenomenon has been previously reported in fluorescently labeled aptamers, where decreases in fluorescence anisotropy were utilized to quantify binding interactions across diverse systems^39–43^. In both cases, the decrease in anisotropy seemed to arise from an increased local flexibility, despite an overall increase in molecular size upon binding. This parallel suggests that the mechanisms governing anisotropy depolarization in aptamer systems may also apply to protein-lipid interactions and other contexts. By demonstrating that our recombinant platform can serve as a precise tool for measuring association with negatively charged membranes, this work greatly expands the scope of fluorescence anisotropy-based sensing, showcasing that a decrease in anisotropy can be a highly efficient biomolecular fluorescence probe. In contrast with a traditional fluorescence anisotropy approach, this technique is not constrained by an upper anisotropy limit (easily reached by large proteins) rendering the free and bound states indistinguishable from one another. Furthermore, our findings show that there is no inherent need for lengthy structural investigations for an “ideal” rigid fluorescent labeling site when designing anisotropy probes. Instead, they demonstrate that previously ignored unstructured protein domains can be purposefully employed for binding measurements. Together, these findings reinforce the potential of fluorescence anisotropy as a precise and adaptable method for probing the governing parameters underlying biomolecular interactions.

While the low salt conditions and high membrane charge concentrations used in this study do not fully replicate the biological milieu, we believe the advantages of a controlled, high-throughput system strongly outweigh these limitations. Furthermore, many of the organelles relevant to our investigation (endosomes, mitochondria, etc.) possess an intrinsically high negative surface charge, making our experimental conditions relevant for probing biologically meaningful interactions. While these conditions may not perfectly capture the full complexity of the intracellular environment, the platform nonetheless provides a controlled system for dissecting lipid specific interactions which can be rescaled to understand broader biophysical principles. Most importantly, the ability of our platform to detect AH sensitivity to certain lipid species in a high throughput manner offers an invaluable new approach to uncovering the peptide code governing AH membrane binding. Discovering inducible amphipathic helix function and the parameters affecting its membrane association can allow us to distinguish between the protein domains responsible for binding from those involved in catalytic function, elucidating fundamental principles that govern membrane-associated biochemical pathways.

The overwhelming complexity of lipid species within organelle membranes is a fundamental hallmark of their biochemical identity, enabling a wide array of targeted protein-membrane interactions. Inducible amphipathic helices are key effectors of these interactions, initiating protein trafficking cascades by sensing specific lipid environments. Each residue of an amphipathic helix is likely solvated by a small number of lipids (on the order of one lipid per residue)^44^, implying that a limited subset of lipids is necessary and sufficient for binding. Our high-throughput platform thus enables a powerful approach to investigate the residue-and peptide-level determinants necessary to detect such lipid environments, establishing a functional framework to probe how variations in individual lipid species can modulate protein recruitment and define the peptide code underlying protein targeting, selectivity, and function.

## Supporting information

Supporting Information, Combined

## Acknowledgements

We would like to thank David M. Jameson for insightful conversations. P.J.C. and A.M. are supported by a grant from the National Institutes of Health (5R35GM150716). D.H.J. and W.Z. are supported by a grant from the National Institutes of Health (5R35GM147333). T.S.U. is supported by a grant from the National Institutes of Health (R01AG072442). Special thanks to Nuria Melisa Morales Garcia and Miranta Kouvari for their artistic contributions to Figure 1 and Figure 5.

## Materials and Methods

### Materials

#### For DNA cloning

DH10B Competent cells were purchased from Thermo Scientific (Waltham, MA, USA). E.Z.N.A.® Plasmid DNA Maxi Kit was purchased from Omega Bio-tek (Norcross, GA, USA). Restriction digestion enzymes NdeI and NcoI-HF, Q5^®^ High-Fidelity DNA Polymerase, NEBuilder^®^ HiFi DNA Assembly Master Mix and Monarch^®^ Spin DNA Gel Extraction Kit were purchased from New England Biolabs (Ipswich, MA, USA). Agarose SFR^TM^ was purchased from VWR (Radnor, PA, USA).

#### For protein expression, purification, and labeling

BL21(DE3) Competent E. coli cells were purchased from New England Biolabs. Sodium chloride (CAS #7647-14-5), Sodium phosphate (CAS #7558-80-7), Imidazole (CAS #288-32-4), 4-(2-hydroxyethyl)-1-piparazineethanesulphonic acid (HEPES) (CAS #75277-39-3), isopropyl-β-D-thiogalactopyranoside (IPTG) (CAS #367-93-1), Tris(2-carboxyethyl)phosphine hydrochloride (TCEP-HCl) (CAS #51805-45-9) were purchased from VWR International. LB Miller broth and LB Miller agar were purchased from IBI Scientific (Dubuque, IA, USA). PureCube 100 Ni-NTA Agarose beads were purchased from Cube Biotech (Monheim, Germany). MilliporeSigma™ Amicon™ centrifugal filter units with 3kDa MW cutoff were purchased from Sigma-Aldrich (St. Louis, MO, USA). Oregon Green™ 488 Maleimide, Alexa Fluor™ 647 C_2_ Maleimide and dimethyl sulfoxide (DMSO, CAS #67-68-5) were purchased from Thermo Fisher Scientific. Isotopically enriched D-Glucose-^13^C_6_ (CAS #110187-42-3) and ammonium-^15^N chloride (CAS #39466-62-1) were purchased from Sigma-Aldrich. All reagents were used without additional purification.

#### For liposomes

1,2-dioleoyl-sn-glycero-3-phosphocholine (DOPC), 1,2-dioleoyl-snglycero-3-phospho-L-serine (DOPS), 1,2-dioleoyl-sn-glycero-3-phosphoethanolamine (DOPE), 2-dipalmitoyl-sn-glycero-3-[(N-(5-amino-1-carboxypentyl)iminodiacetic acid)succinyl] (DGS-NTA(Ni)), 1’,3’-bis[1,2-dioleoyl-sn-glycero-3-phospho]-glycerol (Cardiolipin) and cholesterol (plant-derived) were purchased from Avanti Polar Lipids, Inc (Alabaster, AL, USA). Chloroform (CAS #67-66-3) was purchased from VWR International.

#### For experimental measurements

Corning® 96-well Half Area Black Flat Bottom Polystyrene NBS Microplate were purchased from Corning (Corning, NY, USA). Quartz fluorometer cell cuvettes were purchased from Starna Cells (Atascadero, CA, USA). Colloidal silica (CAS #112926-00-8) was purchased from Electron Microscopy Sciences (Hatfield, PA, USA).

### Methods

#### Plasmids

A master plasmid was designed and produced using a pET28a(+) backbone with an inserted domain encoding our platform (Twist Bioscience, San Francisco, CA, USA). The domain encodes a NdeI enzyme cleavage site, followed by a filler non-encoding region, a NcoI enzyme cleavage site, the sequence encoding for SUMO(20-98) (Uniprot: Q12306) with a mutated cysteine in the C-terminus, a sequence for the TEV cleavage site (ENLYFQS) and a hexa-histidine tag followed by a stop codon. For each peptide of interest, a separate plasmid was cloned by purchasing a linear double stranded DNA fragment from Azenta (South Plainfield, NJ, USA) encoding for the corresponding peptide sequence, followed by the sequence for a GSGS soluble linker region and the SUMO(1-20) domain. The linear DNA fragment was cloned into the master plasmid using NEBuilder^®^ HiFi DNA Assembly. After cloning, plasmids were transformed into DH10b chemically competent cells. DNA was isolated using the E.Z.N.A.® Plasmid DNA Maxi Kit and stored frozen in-20°C. Sequences were confirmed using full plasmid sequencing by Plasmidsaurus (San Francisco, CA, USA). The pNatB (pACYCduet-naa20-naa25) duet plasmid (Plasmid #53613) and the MBP-superTEV (Plasmid #171782) encoding plasmid was acquired from Addgene. Plasmids will be available on Addgene.

#### Protein Expression and Purification

Plasmids were transformed in BL21(DE3)-pNatB^25^ chemically competent cells using standard transformation protocols. A saturated overnight starter culture from a single colony was used to inoculate 1L of LB which was grown at 37℃ to an OD of 0.5 and induced with 1mM of IPTG for 4 hours. Liquid colonies were centrifuged at 4,000*g* for 20 minutes at 4℃ and the pellets were resuspended in Ni-Reaction Buffer: 50mM sodium phosphate @ pH 8, 500 mM NaCl, 10mM Imidazole and sonicated for 10 minutes (2s on, 2s off, 50% amplitude) with a Qsonica Q125 (Qsonica, Newtown, CT, USA). Bacterial lysate was separated by centrifugation at 25,000 g for 30 minutes at 4℃. Proteins were then purified by a 40-minute rotating incubation with NTA-Ni beads. Bound proteins were eluted with 50mM sodium phosphate, 500 mM NaCl, 300mM Imidazole (pH 8) and then buffer exchanged into 50mM Tris, 0.5 mM EDTA, 0.5 mM TCEP (pH 8) using an Amicon centrifugal concentrator filter (3kDa cutoff). TEV protease (Addgene plasmid #171782) was then added and incubated at 37℃ (not shaken) for 1 hour and then overnight at 4℃. The sample was buffer exchanged into Ni-Reaction buffer using an Amicon filter and incubated with NTA-Ni beads. This time the flowthrough was collected since the protein of interest has had its hexa-histidine tag cleaved. The sample was then further purified and buffer exchanged into 20mM Hepes, 150 mM NaCl, 0.5mM TCEP (pH 7.2) by size exclusion chromatography using a Cytiva Superdex 75 Increase 10/300 column with an AKTA Pure system (Cytiva, Marlborough, MA, USA). Peak fractions were collected, mixed, aliquoted, flash frozen and stored in-80℃.

#### Protein Labeling

Oregon Green 488-maleimide and Alexa Fluor 647-maleimide dyes were dissolved in DMSO and stored in-20℃. Labeling of the cysteine residue of each protein construct was performed at 100μM protein concertation with a 5x molar excess of dye in 20 mM Hepes @ pH 7.2, 150mM NaCl, 0.5mM TCEP (pH 7.2) overnight at 4℃. The amount of DMSO in the reaction never exceeded 5%. Unconjugated dye was removed using a Cytiva Superdex 75 Increase 10/300 column with an AKTA pure system. Labeling ratios were measured using UV-vis spectroscopy with a Nanodrop One and ranged from 0.75 to greater than 0.95. Labeled proteins were flash frozen and stored at-80℃.

#### Protein expression ^13^C/^15^N-labeled protein

Plasmids were transformed in BL21(DE3)-pNatB^25^ chemically competent cells using standard transformation protocols. A saturated overnight starter culture from a single colony was used to inoculate 1L of LB which was grown at 37℃ to an OD of 0.7. The culture was then spun down at 3000g for 15 min in sterile bottles, and the resulting pellet was resuspended in 1 L of minimal media supplemented with 2g of ^13^C Glucose and ^15^NH_4_Cl. The resuspended culture was grown at 37 ℃ for 1 hour and then induced with 1mM of IPTG for 4 hours. The purification process was the same as described for unlabeled protein.

#### Preparation of vesicles

Lipid mixtures were prepared by combining the appropriate molar concentrations of lipid aliquots suspended in chloroform stored in-80℃. The lipid mixture was then dried using a ThermoScientific Speedvac SPD130DLX evaporator with a Savant RVT5105 refrigerated vapor trap by centrifuging under vacuum for 3 hours. The dried lipid mixture was dried under vacuum overnight and rehydrated the next day using the appropriate buffer and shaken at 37℃ for 45 minutes. The rehydrated mixture was then freeze-thawed 5 times, alternating between a dry ice ethanol bath and a 42℃ water bath. The solution was then extruded 20 times through a track-etched Whatman Nuclepore Track-etch membrane (Cytiva, Marlborough, MA, USA) of desired pore diameter using a Lipex 10mL Extruder (Evonik, Burnaby, BC, CANADA) lipid extruder. The final vesicle size was confirmed by dynamic light scattering.

Size is reported as the z-average. For cardiolipin-containing vesicles, measurements were acquired within 1 day. In all other cases, measurements were acquired at maximum within 3 days.

#### Tryptophan fluorescence

Tryptophan fluorescence spectra were measured on a Horiba Fluorolog QM Spectrophotometer (Kyoto, Kyoto, Japan). The excitation beam monochromators were set at 280nm with 8nm slit widths. The emission monochromators were set to 300-420 nm with 4nm slit widths. For all tryptophan fluorescence experiments each datapoint corresponds to a sample volume of 2.1 mL under buffer conditions of 20 mM HEPES (pH 7.0) @ 25 °C. To vary the lipid/protein ratio, we alter the lipid concentration in the sample. For the comparison experiments between tryptophan fluorescence and fluorescence anisotropy the protein concentration used in the samples were 125 nM for the Amphiphysin, CHMP4B and Huntingtin protein platforms and 100 nM for the Endophilin-B1 platform. To account for the increasing amount of scattering introduced by high lipid concentrations, we acquire a background fluorescence signal of the lipid sample prior to the addition of protein. Spectra were fit to a linear combination of the fully bound and unbound spectra^32,45,46^:

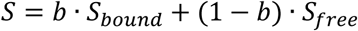

Where b is the molar fraction of the bound protein. The free state spectrum is acquired with protein solely suspended in buffer. We define the fully bound state as the condition in which further increases in lipid concentration no longer alter the observed spectra. Using a minimization algorithm, we determine the b value for every point in between. The b values are then used in nonlinear least square analysis to extract the binding kinetics parameters.

#### Fluorescence Anisotropy

Fluorescence anisotropy measurements were performed on a black 96 well flat bottom half area plate (Corning) using a Biotek Synergy Neo2 plate reader (Agilent, Santa Clara, CA, USA) with an 485/530 polarizing cube for protein samples labeled with Oregon Green 488 (all measurements except Figure S8b) and a 620/680 polarizing cube for protein samples labeled with Alexa Fluor 647 (Figure S8b). 100μL of sample was added to each well, with a final protein concentration between 100-250 nM depending on the experiment (Figure 1b: 125 nM, Figure 2: 250 nM, Figure 3: same as in Tryptophan Fluorescence section, Figure 4: 250 nM). For all experiments the buffering conditions were 20 mM HEPES (pH 7.0). NaCl was included only where indicated (Figure 4c). For each set of measurements where results are compared, the gain is kept constant. To calculate the G factor, we measure the anisotropy of fluorescein sodium salt at 1 nM and adjust G accordingly for the FA to be equal to the known value of 0.027^26^. To extract the bound ratio of each datapoint we use the same method as with the tryptophan fluorescence data.

#### Circular Dichroism Spectroscopy

Circular dichroism measurements were collected on a JASCO J-1100 instrument with a 1 mm path length quartz cuvette (JASCO, Oklahoma City, OK, USA). Scans were taken from 195 – 300nm at a 0.5 nm data pitch with 100 nm/min scan speed. The data integration time was 2 seconds, and each scan was performed twice. The buffer used for each experiment was 20 mM HEPES (pH 7.0) @ 25 °C. Samples of vesicles and protein were allowed to equilibrate for at least 15 minutes before spectra were acquired. Each sample contained 8 μM of protein and was blanked by a buffer solution containing vesicles at the appropriate lipid concentration.

#### Tethered Vesicle Assay

Glass coverslips were cleaned with a modified RCA protocol and passivated with PEG-silane, as described previously with slight modification^47^. Briefly, glass slides were soaked in a concentrated KOH/peroxide bath, followed by a concentrated HCl/peroxide bath each at 80°C for 10 minutes. Once cleaned, the slides were passivated by surface treatment with a 7.5 mg/mL PEG-silane solution in isopropanol. The solution was comprised of 5% biotin-PEG-silane (5000 MW) and 95% of mPEG-silane (5000 MW). All materials for creating the PEG-silane solution were handled in a N_2_ environment. Immediately before passivating the slide, 1% v/v of glacial acetic acid was added to catalyze the reaction between silanes and hydroxyl groups on the cover slip. 50uL of the solution was added to the top surface of the cleaned cover slips, spread out, then placed into an oven at ∼ 70° C for 30 - 60 minutes. The slides were then rinsed under DI water to remove excess PEG-silane and dried under N_2_. These slides were stored dry under air for a maximum of 1 week.

Imaging wells were created by placing 0.8 mm thick silicone sheets (Grace Bio-Labs) with 5mm holes on top of the passivated slide. Each well was hydrated with 6 μg of NeutrAvidin (ThermoFisher Scientific, Waltham, MA, USA) in 30μL of 20mM HEPES (pH 7.0) and incubated for 10 minutes. After incubation, each well was rinsed to remove excess NeutrAvidin. Following this step, each well was rinsed with solution containing vesicles to reach a 1μM lipid concentration. After 10 minutes of incubation, excess vesicles were rinsed from the wells with the same buffer and the protein platform labeled with Oregon Green 488 was added at the appropriate concentration.

#### Confocal Microscopy

Imaging was performed on a laser scanning confocal microscope (Leica Stellaris 5). Two excitation lasers were used: 488 nm and 638 nm for labeled protein and vesicles, respectively. The wavelength detection bandwidths used were 493 – 600 nm and 643 – 748nm for labeled protein and vesicles, respectively. A Leica HC PL APO 63x, 1.4 NA oil-immersion objective was used to acquire images, and the zoom factor was set such that square pixel sizes of 70 nm were obtained. All images were acquired using a scan speed of 400 Hz.

#### Fluorescence anisotropy decay

Fluorescence anisotropy decay measurements were acquired with the DeltaDiode module of the Horiba Fluorolog-QM Spectrophotometer with a DeltaDiode DD-485L 479 nm pulsed excitation laser. Total sample size in all cases was 2.1 mL with buffering conditions of 20 mM HEPES (pH 7.0). Protein concentration was 25 nM, labeled with Oregon Green 488, while vesicle concentrations were 25 μM when present. Prior to each measurement, proper alignment of the excitation laser pulse and the emission polarizer was tested using an optically dilute colloidal silica solution. We considered the beams to be fully orthogonal if the polarization value was found to be 0.97 or greater. Parallel and perpendicular intensity emission responses were measured in sequence. Measuring time was set to be 5 minutes for each. After each measurement, we confirmed that the sample photobleaching was not significant (<2%). The emission spectrophotometer was set at 530 nm with 8nm slit widths. Laser excitation pulse frequency was set at 500 kHz. Time binning was set at a window of 100ns with 1024 total bins. Instrument response function for the parallel and magic polarizer angles were measured using a colloidal silica solution of 0.0003% for one minute in each polarization direction. Analysis of the decays was performed in accordance with prior literature^37^. Briefly, anisotropy decay models were each convolved with the corresponding instrument response functions and joint fitting of the parallel and perpendicular intensities was performed using a custom python code of nonlinear least square fitting (via the lmfit wrapper module of python’s SciPy library) using well-defined weights based on single photon counting statistics. Limiting values for each free parameter were set based on a physical understanding of the expected motion of the fluorophore in each case to avoid searching through the whole parameter phase space. For the double rotational correlation model with local hindered motion, the fluorescence lifetime (τ) was limited to values between 3 and 5 ns, the fast rotational correlation time (θ_F_) was limited between 0.1 and 2 ns, the slow rotational correlation time (θ_P_) was limited between 2 and 10 ns for the case of the free protein and between 2 and 10000 ns for the case of the bound protein, the time zero anisotropy (r_0_) was limited between 0.1 and 0.4, the α parameter was limited between 0 and 1 and finally I_0_ was set to be between 10^3^ and 10^6^. For the single hindered rotational correlation model, the fluorescent lifetime (τ) was limited to values between 3 and 5 ns, the fast rotational correlation time (θ_F_) was limited between 0.1 and 2 ns, the time zero anisotropy (r_0_) was limited between 0 and 0.4, the long-time anisotropy r_∞_ was limited between 0 and 0.1 and finally I_0_ was set to be between 10^3^ and 10^9^.

#### NMR experiments

For backbone assignments, ^13^C/^15^N-labeled protein samples with volumes of 320 μL were prepared at concentrations of 0.9 mM in 50 mM MOPS (pH 7.0), 3 mM TCEP and 6% D_2_O solution (Figure S13). In the presence of vesicles (Figure S14), analogous samples were prepared containing 0.5 mM protein and 25 mM of vesicles with composition DOPC/DOPS/DGS-NTA(Ni): 50/40/10. Backbone assignments were achieved using HNCA, HNCO, HNCACB, CBCA(CO)NH and NOESY-HSQC (t_m_ =100 ms) experiments acquired on a Bruker Avance 700 (Bruker, Billerica, MA, USA) spectrometer at 35 °C. Data were processed and analyzed with the nmrPipe package and CARA^48^. The chemical shifts of untagged and tagged protein have been deposited to the BMRB database with accession codes 53091 and 53092, respectively. In the presence of vesicles, backbone assignments remained virtually unchanged (Fig. S14). Combined ^1^H^N^-^15^N chemical shift differences between untagged and tagged protein were calculated according to:

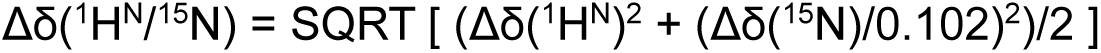

#### Binding kinetics

To analyze inducible amphipathic helix peptide binding with membranes we considered the interaction as a partitioning equilibrium between the water/buffer phase and the lipid bilayer phase as first introduced by White et al^49^ and as applied by Robustelli and Baumgart^13^.

Briefly the partition coefficient K_p_ is given by

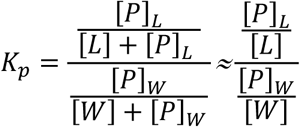

Where [L] is the lipid concentration, [W] is the water concentration and [P]_L_ and [P]_W_ are the protein concentrations in the lipid bilayer and in the buffer solution respectively, with the total amount of protein [P]_t_=[P]_L_+[P]_W_, which allowed us to find the fraction of the protein in the bilayer:

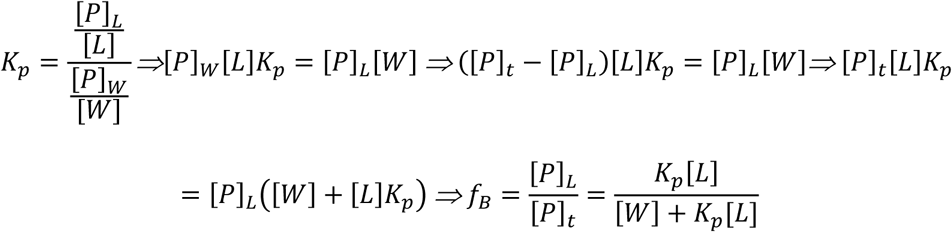

For the high analysis shown in Figure 2, we treated the fluorescence anisotropy signal of an intermediate bound state as a linear combination of the FA of the fully bound (S_B_) and free (S_F_) states:

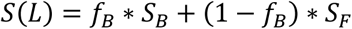

We acquired fluorescence anisotropy data (S(L)) for different [L] values and determined the K_p_ by fitting the following equation:

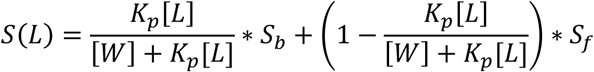

using two fitting parameters K_p_ and S_B_. The water concentration was set to be [W]=55.5 M, with S_f_ determined for [L]=0.

For the comparison measurements between fluorescence anisotropy and tryptophan fluorescence shown in Figure 3, we transformed data acquired with each technique into bound ratio data as described in the “Tryptophan Fluorescence” Methods section. Then, the bound ratio data were fitted using the above equation for S(L), setting S**_F_**=0 with two fitting parameters K_p_ and S_B_. Since we assume a single bound state and, in all cases, we have either reached or are very close to the plateau of the binding curve, the resulting S_B_ is essentially a scaling parameter proportional to the difference in signal between the free and bound states. As such, we re-normalized the resulting fitted curve and bound ratio data to 1 by dividing by the fitted value of S_B_, considering error propagation for the data point error bars.

To analyze the fluorescence anisotropy data from the His_6_-SUMO construct shown in Figure 4, we used the depletion model, treating the protein as a ligand binding to independent binding sites, that are not always greater than the number of ligands in the sample^32^. The bound ratio was then calculated as:

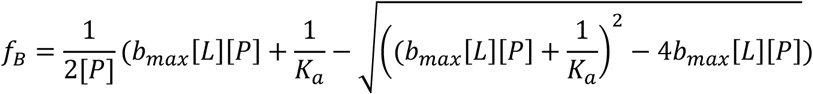

To fit the data, we again treated the fluorescence anisotropy signal of an intermediate bound state as a linear combination of the FA of the fully bound (S_B_) and free (S_F_) states:

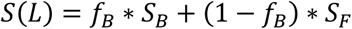

The binding data was fitted using nonlinear least square analysis with two fitting parameters, K_a_ and b_max_.

