## Supporting Information, Combined for "Recombinant protein platform for high-throughput investigation of peptide-liposome interactions via fluorescence anisotropy depolarization"

[1]Department of Physics and Astronomy, University of Southern California, Los Angeles, California, USA. [2]Department of Physiology and Neuroscience, Zilkha Neurogenetic Institute, Keck School of Medicine, University of Southern California, Los Angeles, California, USA. [3]Mork Family Department of Chemical Engineering and Materials Science, University of Southern California, Los Angeles, California, USA. [4]Department of Chemistry, University of Southern California, Los Angeles, California, USA. [5]Alfred E. Mann Department of Biomedical Engineering, University of Southern California, Los Angeles, California, USA.

*Corresponding Author

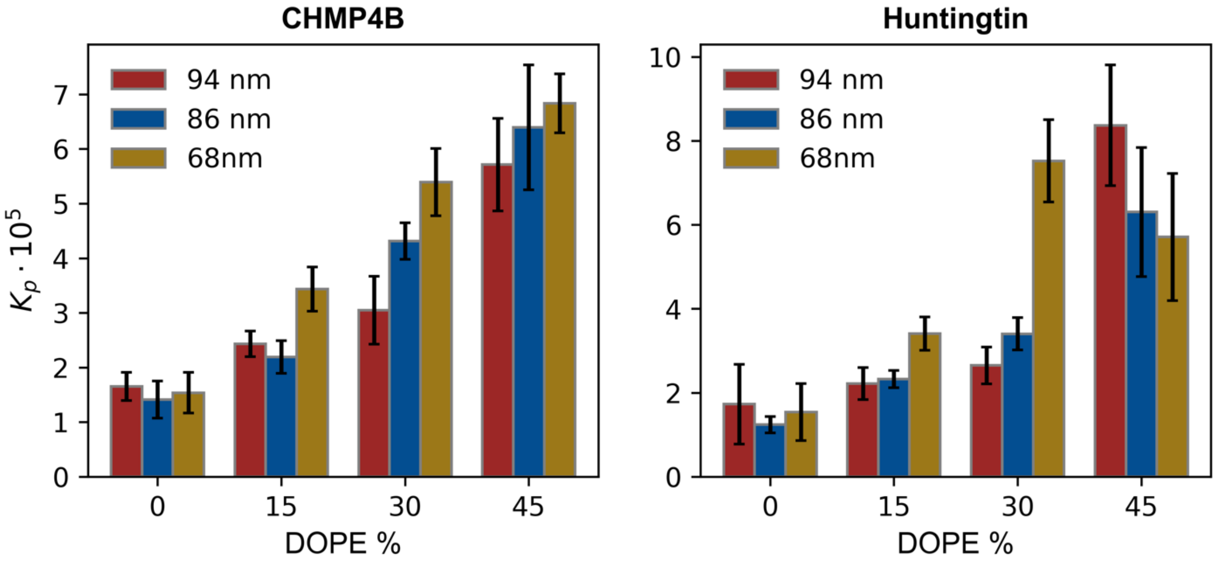


**Figure S1 | Fitted partition coefficient values from fluorescence anisotropy data of Figure 2**. Error bars represent standard errors of each fit. Vesicles composition in each case is DOPC/DOPS/DOPE : (70-X)/30/X where X is the percentage shown on the x-axis. Legend shows measured average vesicle diameter.

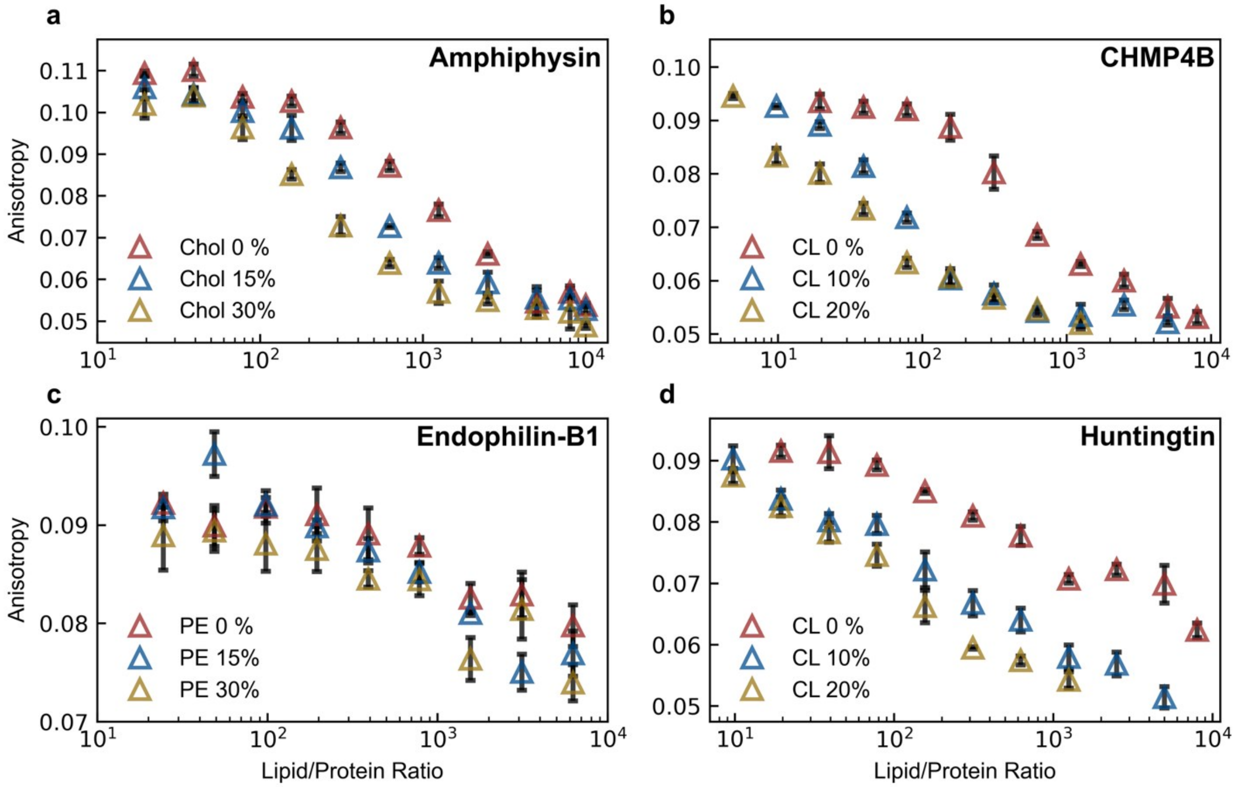


**Figure S2 | Fluorescence anisotropy measurements of the 4 peptide constructs shown in Figure 3**. Each platform was tested against a different array of liposome compositions as described in Fig 3. Data points show mean of 4 replicate measurements for each lipid/protein ratio, with standard deviations plotted (black error bars).

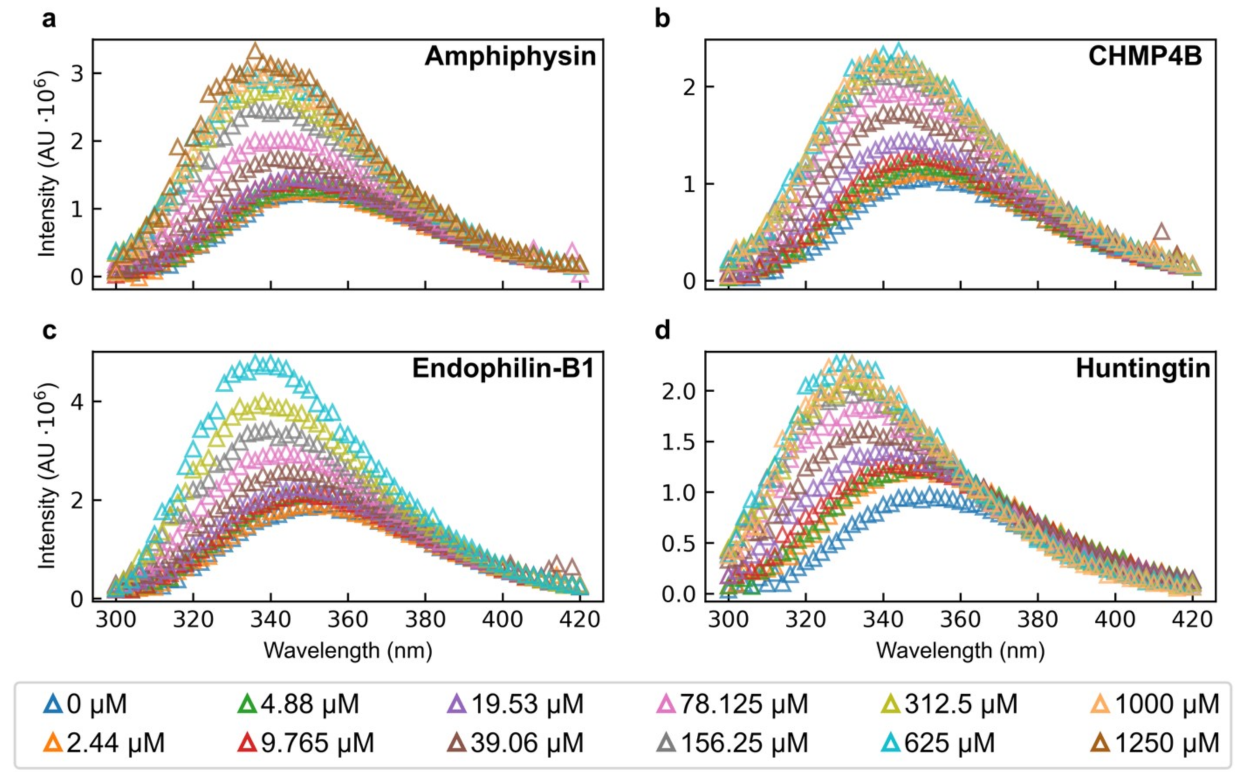


**Figure S3 | Representative tryptophan fluorescence measurement spectra from data shown in Figure 3**. Tryptophan fluorescence spectra of our protein platforms including inducible the amphipathic helices of **(a)** Amphiphysin (AA: 1-25, F9W, 125 nM), **(b)** Endophilin-B1 (AA: 1-33, F18W, 100 nM), **(c)** Huntingtin (AA: 1-17, F11W, 125 nM) and **(d)** CHMP4B (AA: 1-19, F8W, 125 nM) measured against varying concentrations (legend) of DOPC/DOPS:70/30 liposomes with size distribution averages of ~70 nm.

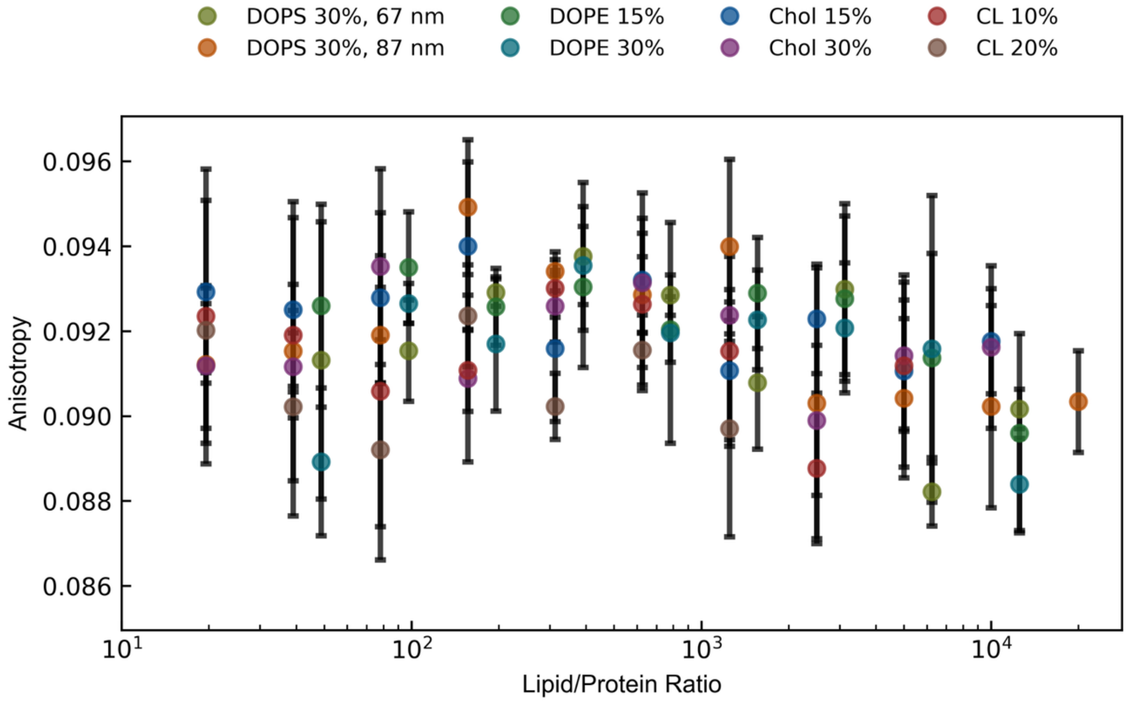


**Figure S*4* | Without conjugated peptides, our platform is inert to lipid vesicles**. Fluorescence anisotropy measurements of fluorescently-labeled recombinant platform with no conjugated peptide with varying liposome compositions. In all cases, vesicles were measured to be ~70 nm in diameter unless otherwise noted, composed of DOPC/DOPS/variable lipid (70-X)/30/X. Variable lipids were DOPE, cardiolipin (CL) and cholesterol (Chol). Protein concentrations were 100-250 nM depending on experiment. Data points show mean of 4 replicate measurements for each lipid/protein ratio, with standard deviations plotted (black error bars).

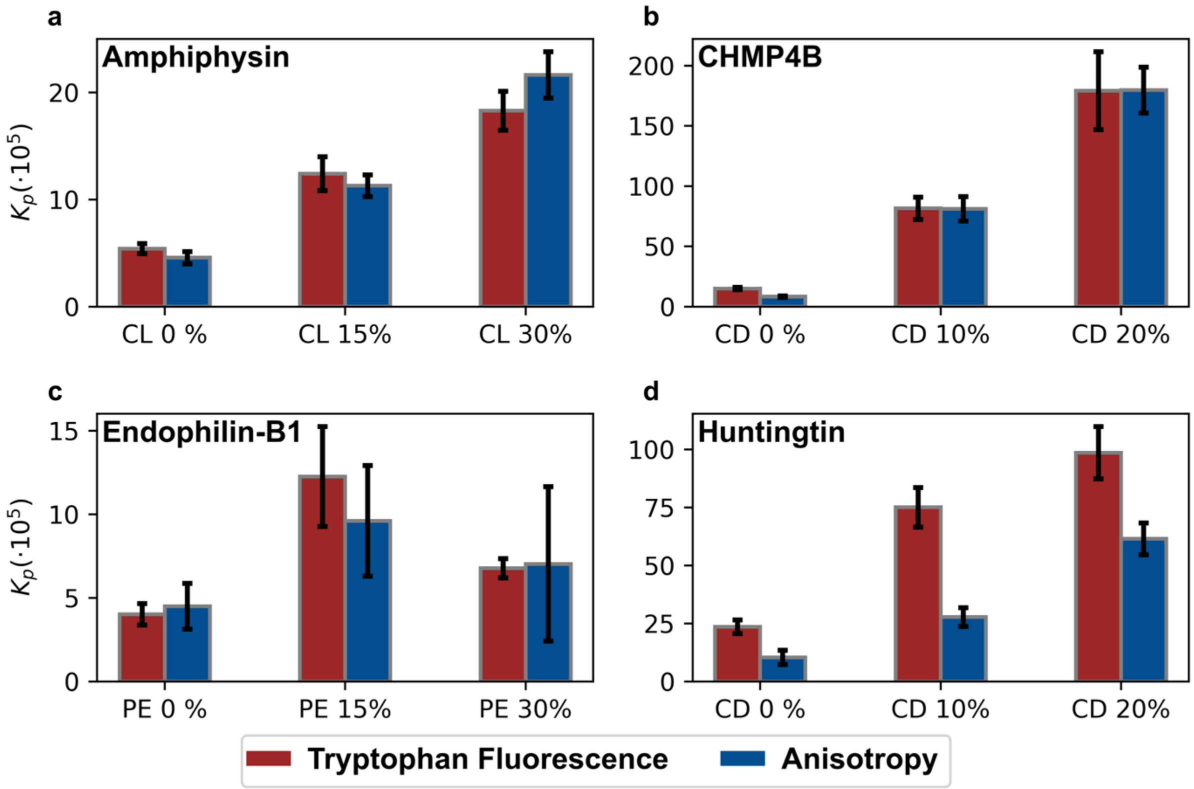


**Figure S5 | Fitted partition coefficient values from Figure 3**. Error bars represent the standard errors of each fit.

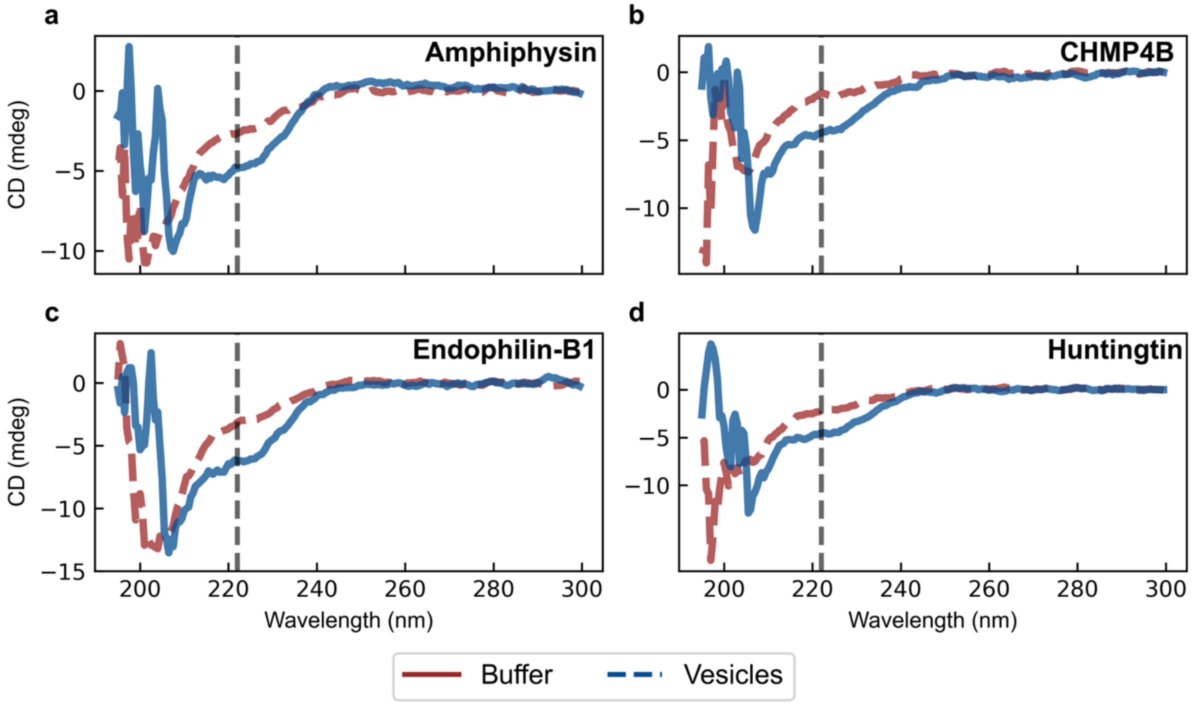


**Figure S6 | Peptides transition into amphipathic helices upon interacting with liposomes.** Circular dichroism spectra of our protein platforms (8 μM) including the inducible amphipathic helices of **(a)** Amphiphysin (AA: 1-25, F9W), **(b)** CHMP4B (AA: 1-19, F8W), **(c)** Endophilin-B1 AA: 1-33, F18W) and **(d)** Huntingtin (AA: 1-17, F11W) measured against 4 mM of DOPC/DOPS:70/30 vesicles with size distribution averages of ~68 nm. Characteristic decrease in CD signal at 222 nm (black dashed line) indicates the formation of an amphipathic helix.

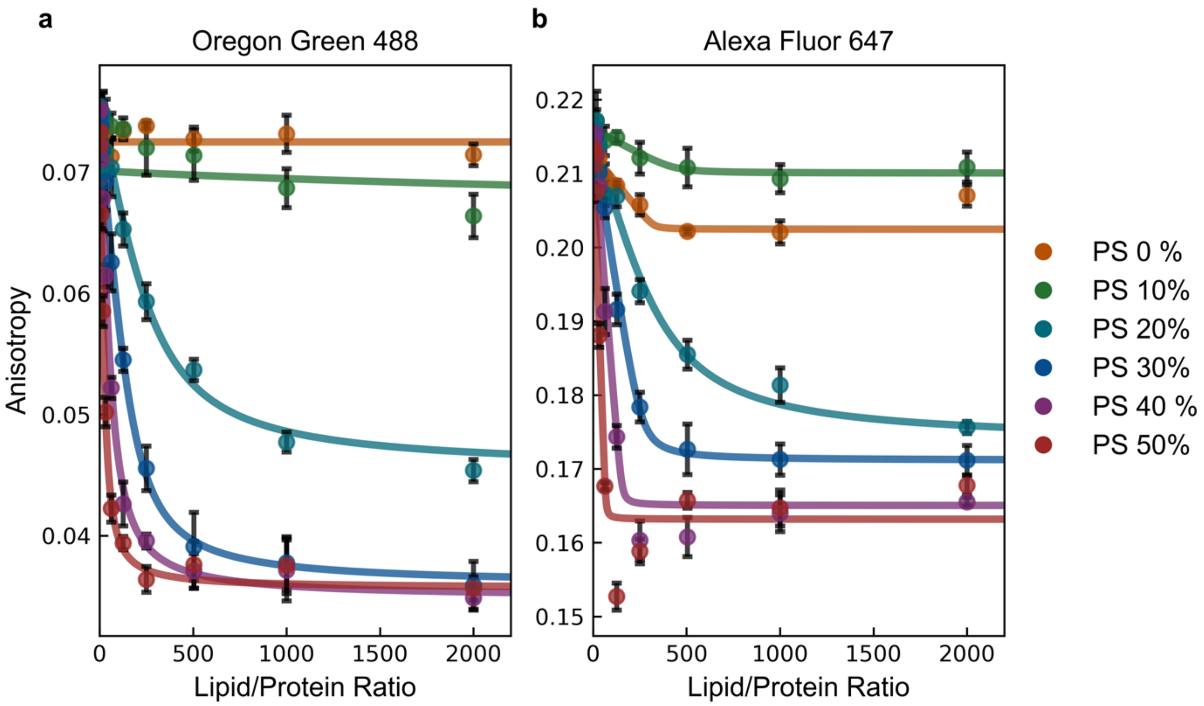


**Figure S7 | The decrease in fluorescence anisotropy is independent of fluorophore identity.** Fluorescence anisotropy measurements of differently fluorescently-tagged hexahistidine model platforms (@ 250 nM concentration) binding to similarly composed nickel-chelating lipid containing vesicles with varying amounts of DOPS membrane content (DOPC/DOPS/DGS-NTA(Ni) = 90-X/X/10).

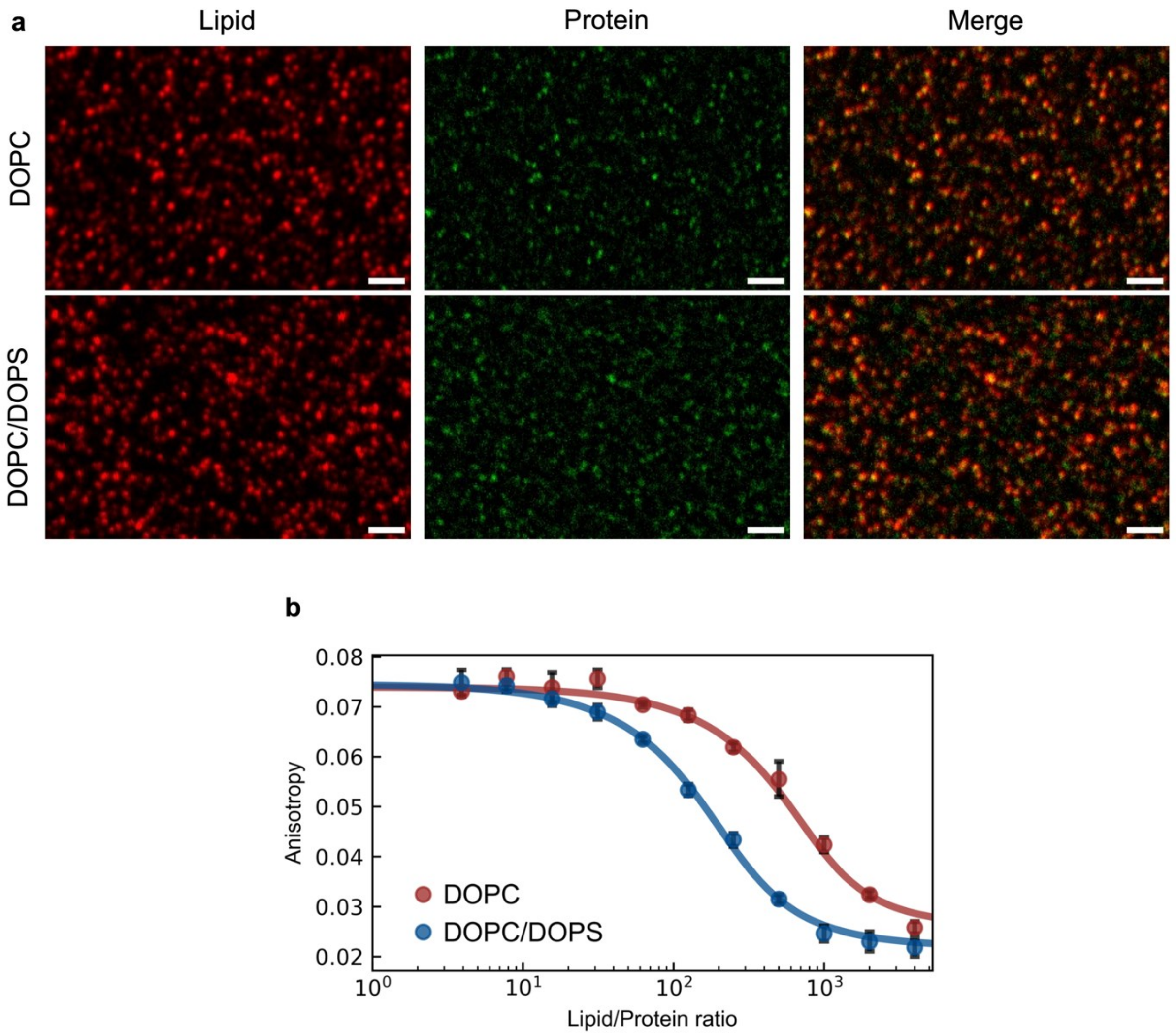


**Figure S8 | Confocal microscopy images confirm binding of hexahistidine model platform to liposomes**. **(a)** Confocal microscopy images from tethered vesicle assay reveal binding of the fluorescently-tagged hexahistidine model platform to vesicles compositions of DOPC/DGS-NTA(Ni)/Biotinyl/DPPE-Atto647 = 87.5/10/2/0.5 vesicles and DOPC/DOPS/DGS-NTA(Ni)/Biotinyl/DPPE-Atto647 = 47.5/40/10/2/0.5. **(b)** Fluorescence anisotropy response of the fluorescently-tagged hexahistidine model platform with the same vesicles as above. A decrease in anisotropy is now present even with non charged vesicles.

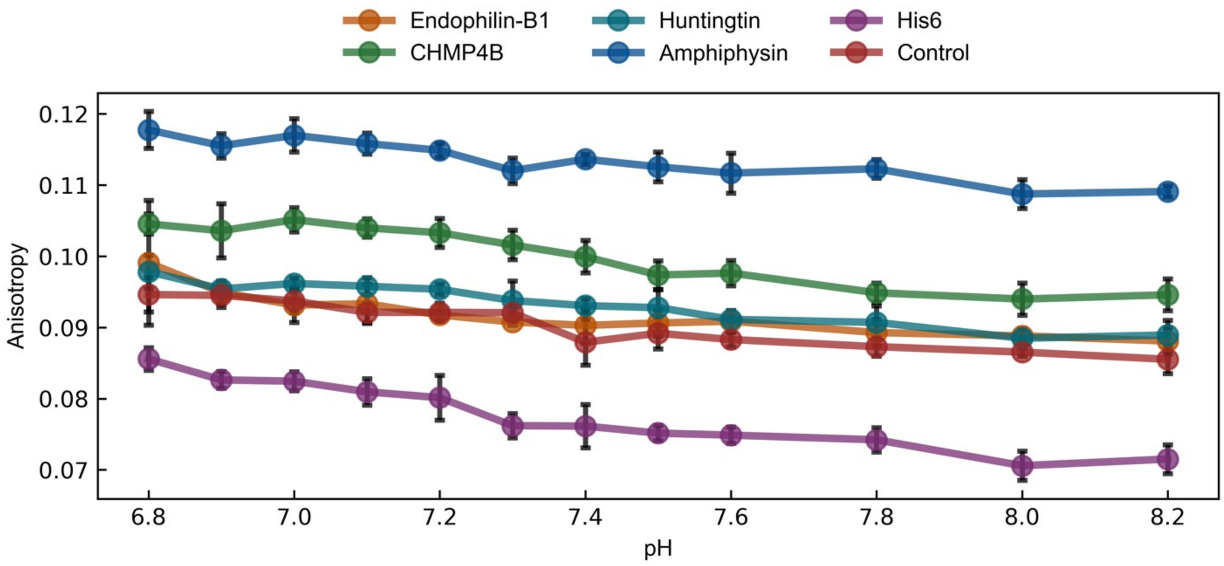


**Figure S9 | Fluorescence anisotropy response of inducible amphipathic helix peptide and control platforms in varying pH**. Fluorescence anisotropy measurements against a pH gradient (20 mM HEPES) were taken for all of all 6 recombinant protein platforms (250 nM) used in this paper. “Control: referes to the fluorescently-labeled recombinant platform with no peptide on its N-terminus as in Figure S2. Data points show mean of 4 replicate measurements for each lipid/protein ratio, with standard deviations plotted (black error bars).

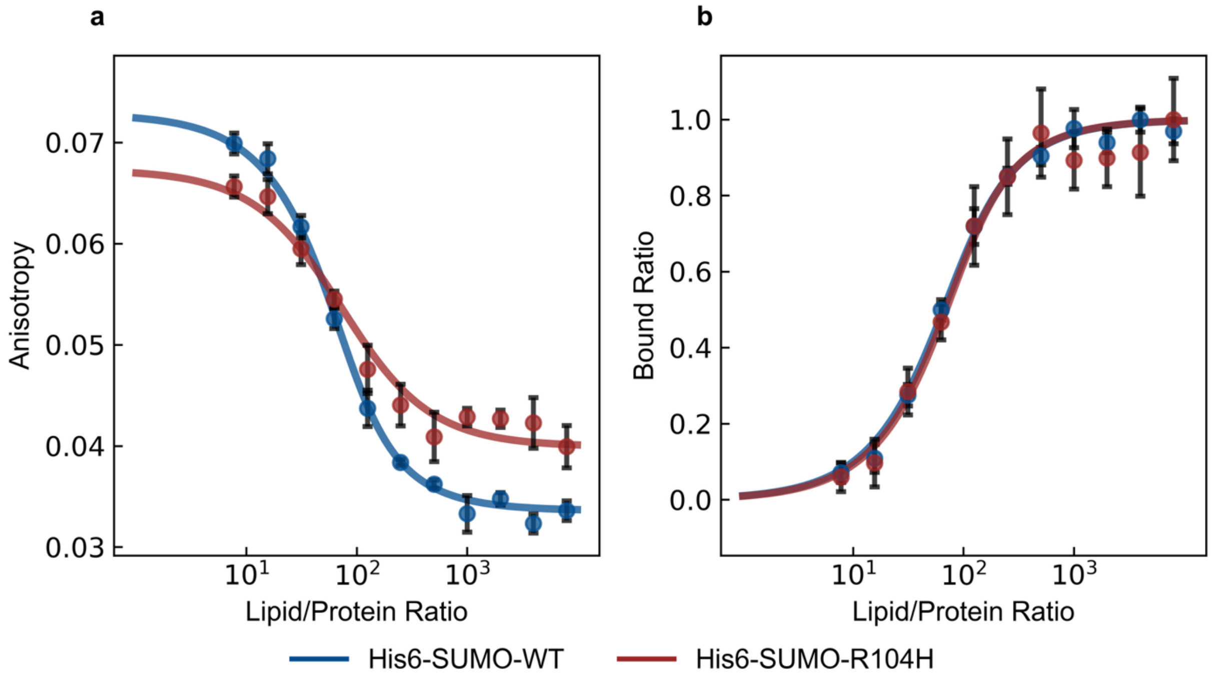


**Figure S10 | Binding response of the fluorescently-tagged hexaxistidine model platform and the mutated R104H model platform to nickel-chelating containing lipid vesicles**. **(a)** Raw fluorescence anisotropy data from mutated model R104H hexahistidine platform (250 nM) binding to vesicles (with a composition of DOPC/DOPS/DGS-NTA(Ni) = 50/40/10 and an average diameter of ~95 nm) and **(b)** transformed data into bound ratio curves as described in Methods. Error bars represent standard deviation from 4 replicate microplate wells.

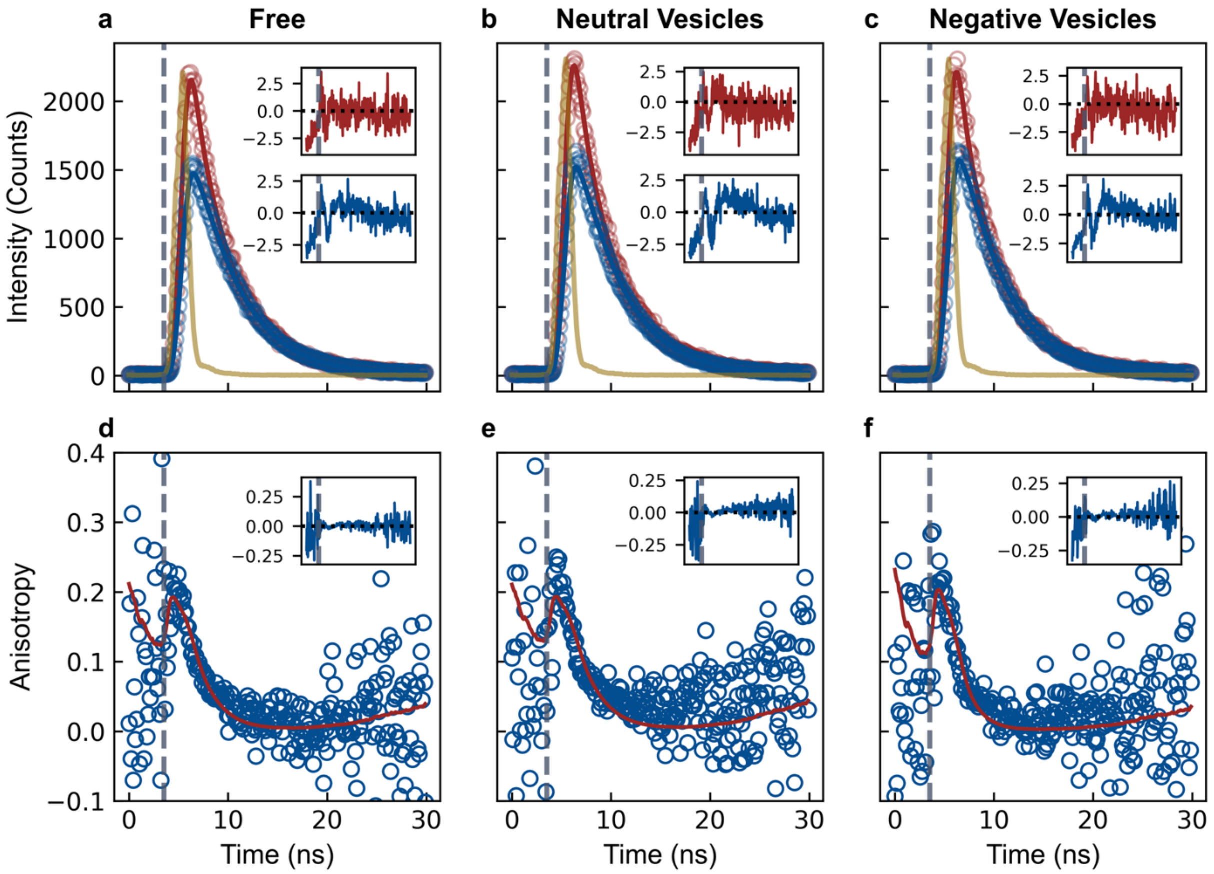


**Figure S11 | Anisotropy decay data and fits with single rotational correlation lifetime model**. Parallel (blue) and perpendicular (red) intensity decays and corresponding fits of the fluorescently-tagged hexahistidine model platform (25 nM) **(a)** free in solution **(b)** bound to neutral vesicles (DOPC/DGS-NTA(Ni) = 90/10, 25 μM) and **(c)** bound to negatively charged vesicles (DOPC/DOPS/DGS-NTA(Ni) = 50/40/10, 25 μM). **(d-f)** Clalculated anisotropy decays and fits from (a-c) respectively. Insets show residuals of each fit. Vertical dashed grey line indicated start of pulse. Yellow curve indicates the instrument response function.

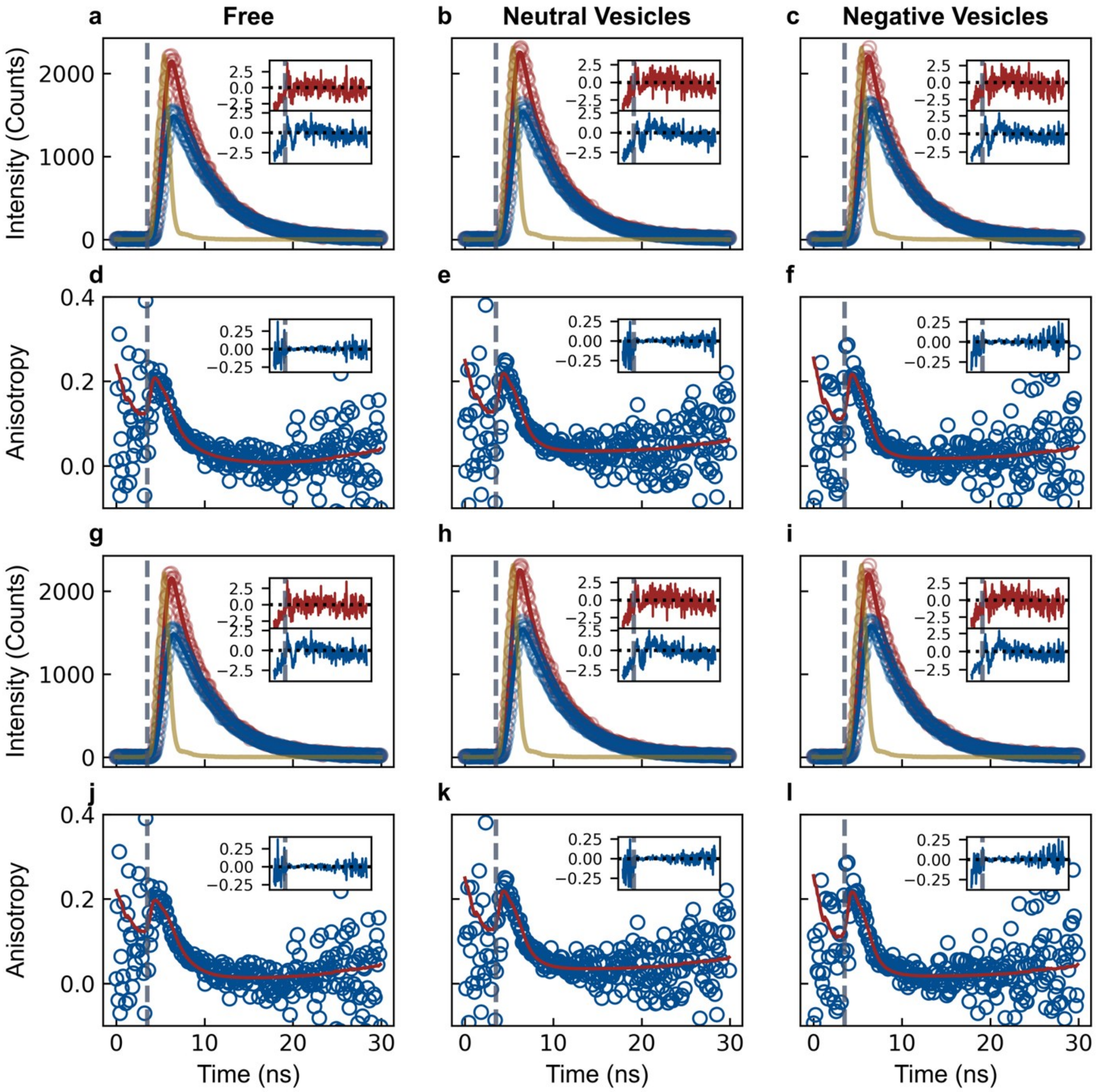


**Figure S12 | Anisotropy decay data of fluorescently-tagged hexahistidine model platfrom fitted with 2 different models**. Anisotropy decay data from Figure S11 fitted with 2 different models shown in 3 columns (**Free**: protein free in solution, **Neutral Vesices**: protein bound to vesicles comprised of DOPC/DGS-NTA(Ni) = 90/10, **Negative Vesicles**: bound to negatively charged vesicles (DOPC/DOPS/DGS-NTA(Ni) = 50/40/10). **(a-c)** Parallel (blue) and perpendicular (red) intensity decays and corresponding fits using the two-state hindered anisotropy decay model. **(d-f)** Clalculated anisotropy decays and fits from (a-c) respectively. **(g-i)** Parallel (blue) and perpendicular (red) intensity decays and corresponding fits using reduced two-state model (hindered rotational diffusion) model. **(j-l)** Clalculated anisotropy decays and fits from (g-i) respectively. Insets show residuals of each fit. Vertical dashed grey line indicated start of pulse. Yellow curve indicates the instrument response function.

**Hindered Rotational Diffusion in Membranes**

In order to effectively evaluate the fitted fluorescence anisotropy decay data and produce a more comprehensive geometric interpretation of the phenomena detected, we employ an interpretation of the hindered rotation describing the molecule as rod-like in an infinite square-well potential so that its rotation is unhindered until a certain angle $\theta_C$ is reached[1–3]. In this model the limiting and time zero anisotropy are related to the cone angle:

$$\frac{r_\infty}{r_0} = \left(\frac{1}{2} cos\theta_C (1 + cos\theta_C)\right)^2$$

with completely unhindered motion for $\theta_C$= 90°. This approach can be expanded to the dual rotational diffusion with hindered segmental mobility model by considering the relationship between a and $r_\infty$: $r_\infty = r_0(1 - a)$ and hence extract an estimated angle for the cone of motion for the local hindered diffusion of the fluorophore. For the free state we calculate a $\theta_C$≈40±10° whereas for the bound state on neutral and negative vesicles it is $\theta_C$≈61±2° and $\theta_C$≈69±2° respectively.

**Table S1 | Fitted parameter values from anisotropy model fits.** 1r: single rotational correlation lfietime model, 2r: two-state hindered anisotropy decay model, 2r-H-red: reduced two-state model (hindered rotational diffusion) model

| | $I_0$ (AU) | $\tau$ (ns) | $r_0$ | $\alpha$ | $\theta_F$ (ns) | $\theta_P$ (ns) | $r_{inf}$ | $\theta_{cone}$ (°) |
|---|---|---|---|---|---|---|---|---|
| **1r Free** | 7290±30 | 4.25±0.01 | 0.212±0.007 | - | - | 1.9±0.1 | - | - |
| **1r Bound Neutral** | 7610±40 | 4.18±0.01 | 0.211±0.007 | - | - | 2.2±1.1 | - | - |
| **1r Bound Negative** | 7660±39 | 4.17±0.01 | 0.23±0.01 | - | - | 1.30±0.09 | - | - |
| **2r Free** | 7290±30 | 4.24±0.01 | 0.24±0.02 | 0.6±0.3 | 1.1±0.7 | 3±1 | - | 40±10 |
| **2r Bound Neutral** | 7630±30 | 4.16±0.01 | 0.25±0.01 | 0.87±0.03 | 1.1±0.1 | ~∞ | - | 61±3 |
| **2r-H Bound Negative** | 7670±30 | 4.15±0.01 | 0.25±0.01 | 0.94±0.03 | 0.9±0.1 | ~∞ | - | 69±4 |
| **2r-H-red Bound Neutral** | 7630±30 | 4.16±0.01 | 0.25±0.01 | - | 1.1±0.1 | - | 0.033±0.004 | 61±2 |
| **2r-H-red Bound Negative** | 7670±30 | 4.15±0.01 | 0.25±0.01 | - | 0.93±0.09 | - | 0.016±0.004 | 69±2 |

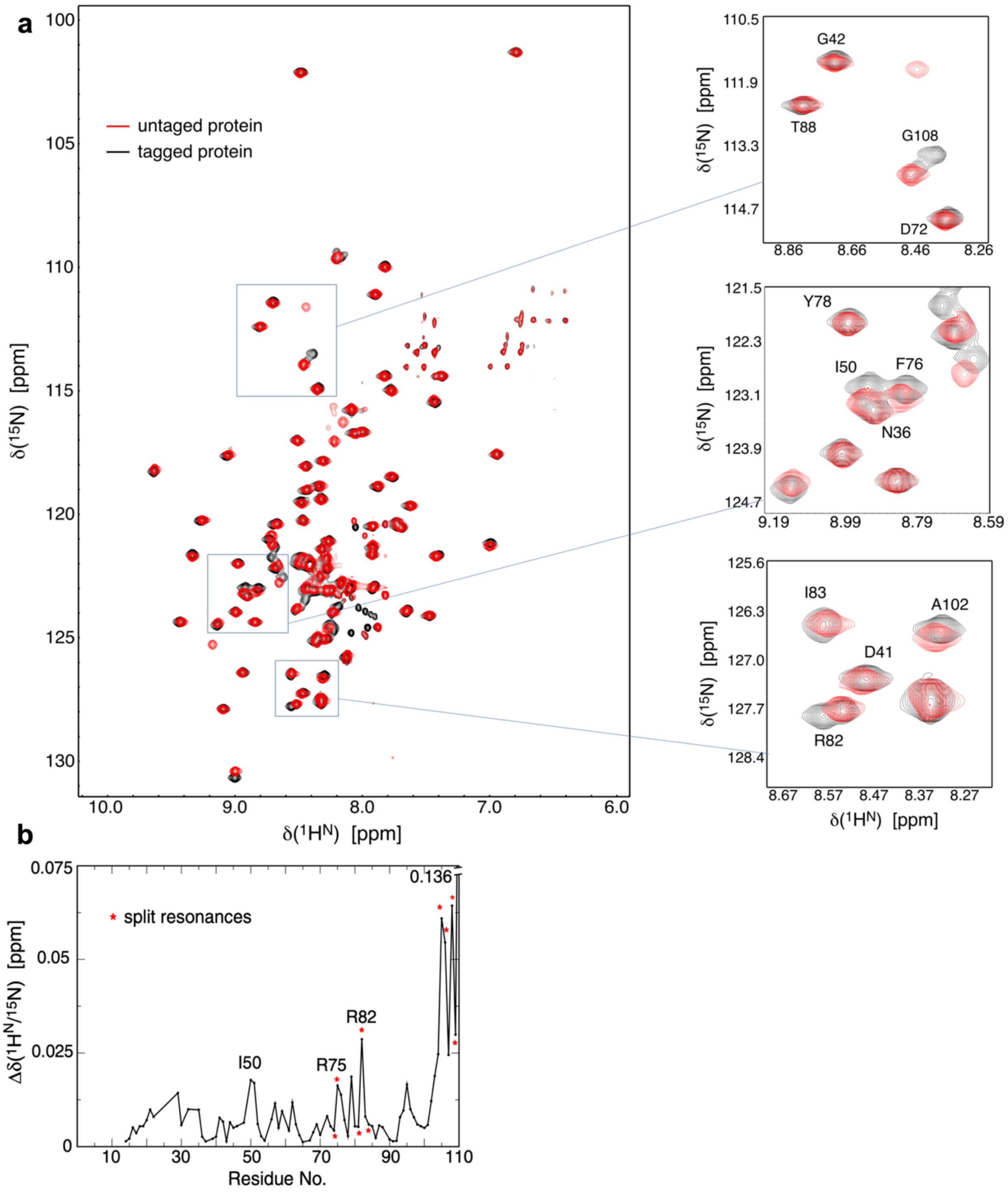


**Figure S13 | Comparison of $^{1}H^{N}$-$^{15}N$ correlation spectra of fluorescently-tagged and untagged hexahistidine model platform. (a)** For C-terminal residues shifted peaks and resonance splitting were prevalent. Nonetheless, this pattern also occurred within the SUMO domain as indicated. **(b)** Chemical shift difference between Oregon Green 488 tagged and untagged hexahistidine model platform. Some residues exhibited doubled resonances. For such split peaks, the most intense signal was selected.

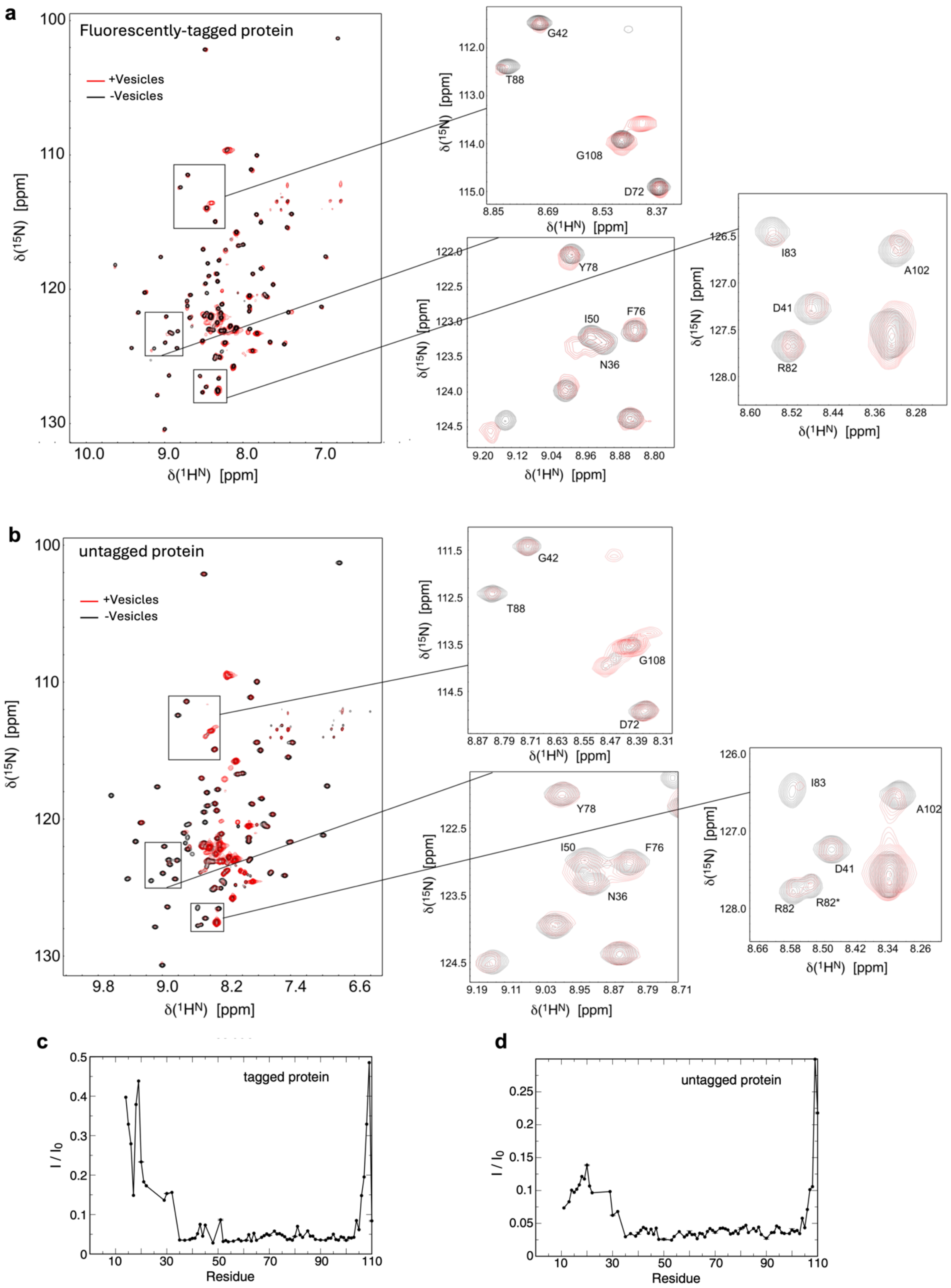

a
Fluorescently-tagged protein
+Vesicles
-Vesicles
δ(15N) [ppm]
δ(1HN) [ppm]
G42
T88
G108
D72
Y78
I50
F76
N36
I83
A102
D41
R82
b
untagged protein
+Vesicles
-Vesicles
δ(15N) [ppm]
δ(1HN) [ppm]
G42
T88
G108
D72
Y78
I50
F76
N36
I83
A102
D41
R82
R82*
c
tagged protein
I / I0
Residue
d
untagged protein
I / I0
Residue

**Figure S14 | Comparison of $^{1}H^{N}$-$^{15}N$ correlation spectra of free and vesicle-bound hexahistidine model platform. (a)** Correlation spectra of fluorescently-tagged and **(b)** untagged hexahistidine model platform when free in solution vs when partially bound to vesicles comprised of DOPC/DOPS/DGS-NTA(Ni) = 50/40/10. In the presence of vesicles, spectral positions of the protein platform remained close to the vesicle-free state with reduced signal intensities, indicating predominantly slow exchange kinetics between free and vesicle-bound protein. Because of the large overall particles size, resonances of vesicle-bound protein are unobservable. **(c, d)** Residual $^{1}H^{N}$-$^{15}N$ cross-peak intensities of fluorescently-tagged and untagged hexahistidine model platform in the presence of vesicles. For both platforms, the same three regions of signal reductions were observed illustrating similar degrees of domain flexibility.
